# A tonsil organoid model reveals Epstein-Barr virus infected germinal center B cell states during primary infection

**DOI:** 10.64898/2026.01.15.699764

**Authors:** Mahina Tabassum Mitul, Yizhe Sun, Timothy B. Yates, Suhas Sureshchandra, Jenna M. Kastenschmidt, Andrew M. Sorn, Zachary W. Wagoner, Erika M Joloya, Arjun Nair, Allyssa Daugherty, Naresha Saligrama, Gurpreet Ahuja, Qui Zhong, Douglas Trask, Leah C. Kottyan, Matthew T. Weirauch, Carmy Forney, Benjamin E. Gewurz, Lisa E. Wagar

## Abstract

Epstein-Barr virus (EBV) colonizes secondary lymphoid tissues to establish persistent infection and is strongly associated with malignancy and autoimmunity. Our understanding of EBV infection biology is hindered by a lack of models that capture infected B cell activity in the lymphoid tissue microenvironment. We therefore developed an EBV human tonsil organoid model to evaluate key B cell states and antiviral responses, including after primary infection. EBV promoted B cell differentiation into germinal center (GC)-like phenotypes and transcriptomic analyses highlighted numerous B cell transcriptional programs unique to EBV-infected cells. B cell receptor repertoire analysis revealed that most EBV-infected B cells underwent class switching but only rarely participated in somatic hypermutation. CD4 T cells, highly activated by organoid infection, limited EBV^+^ B cell outgrowth. Our findings demonstrate human tonsil organoids as a physiologically relevant model to investigate key aspects of EBV immunity and pathogenesis.

## Main

Epstein-Barr virus (EBV) is a B lymphotropic, gamma-herpesvirus^1^. EBV causes a range of B cell lymphomas, T and NK lymphomas, gastric and nasopharyngeal carcinomas, and autoimmune diseases^2–4^. Over 95% of adults worldwide are chronically infected^1^. Most primary infections are asymptomatic and occur in early childhood. However, when initial infection occurs in adolescence or in early adulthood, a third to a half of individuals develop infectious mononucleosis (IM), a syndrome characterized by massive lymphocyte and natural killer cell activation^5^. Waldeyer’s ring, tissue including tonsils, is the primary site of EBV infection, and B cells are the dominant cell target of persistent infection^6^. Like other herpesviruses, EBV exhibits a biphasic life cycle, consisting of latent and lytic phases, each defined by distinct viral gene and protein expression patterns^7^. After resolution of primary infection, EBV establishes latency in a small pool of memory B cells^8^, from which it sporadically reactivates upon plasma cell differentiation to support transmission^9^.

There has been great interest and effort to understand EBV’s complex viral life cycle. According to the prevailing “germinal center” model^6^, EBV is thought to initially infect naive B cells and trigger their entry into germinal centers (GCs), specialized secondary lymphoid regions of B cell proliferation, somatic hypermutation and class-switch recombination (CSR). Infected B cells are posited to undergo sequential changes in viral latency gene expression. EBV latency programs are comprised of one to eight viral protein-encoding genes or non-coding RNAs (ncRNAs)^10–12^ and are variably immunogenic depending on the number, type, and functions of the expressed genes. Due to the asymptomatic nature of most primary infections, latency programs have been mostly based on studies of *in vitro* generated lymphoblastoid cells (LCLs), *ex vivo* analysis of EBV-induced lymphomas and IM^13,14^ and PCR analyses of bulk peripheral blood or tonsillar cells^6,15,16^. *In vivo*, EBV^+^ B cells have been rarely identified in GCs expressing canonical GC markers, including *BCL6* and *AID*^17^. Although these infected GC B cells proliferate, they are few in number and display restricted expansion compared to uninfected GC B cells^18^. EBV^+^ GC B cells are hypothesized to differentiate into memory B cells, the reservoir for lifelong EBV infection. EBV+ memory B cells can be class switched and can show evidence of somatic hypermutation (SHM)^19^, but EBV^+^ tonsil GC B cells may not participate in the GC response^20^. A recent *in vitro* study of EBV^+^ peripheral blood B cells showed that they transcriptionally resemble GC B cells^21^. However, the lack of a tractable human secondary lymphoid model system has left it unclear whether infected B cells actively participate in a GC response during asymptomatic acute or chronic infection.

EBV-associated diseases remain rare, indicating host immunity effectively controls viral pathogenesis in most people. Early viral control is mediated by NK cells and EBV-specific CD4 and CD8 T cells^22–25^. EBV-specific memory T cells are polyfunctional and maintain a virus-host balance in the chronically infected host^26,27^. Antigen-specific CD8 T cells expand during IM and are critical for controlling EBV^+^ B cell proliferation, while CD4 T cells clearly limit EBV pathogenesis as underscored by increased EBV-associated lymphoproliferative disease in HIV-infected individuals^28^.

Despite numerous efforts, no effective vaccine exists to block primary infection or protect against EBV-associated diseases. Use of animal models to understand EBV infection biology has been a challenge because EBV does not infect mice^29^. Although humanized mouse models have been tremendously helpful in understanding the foundational biology of EBV pathogenesis^30,31^, they have limitations related to lymphoid tissue architecture and GC and memory B cell development^30,32,33^. Due to its asymptomatic nature, it is rarely possible to follow the natural course of the immune response to primary EBV infection in humans *in vivo*. To overcome these challenges and complement existing models, we used a human tonsil organoid model^34–38^ to track the early dynamics of EBV infection in tonsillar B cells along with concurrent cellular responses.

### Results

#### Establishing and profiling EBV infection in human tonsil organoids

Tonsil organoids were generated from tonsillectomy patient samples (**Extended Data Table 1**) and infected with the fully transforming Akata EBV strain^39^, enabling EBV^+^ B cell tracking. EBV titration was performed to establish optimal viral doses (**Fig. 1a; Extended Data Fig. 1a** for representative gating). GFP^+^ B cells were rare at early time points (median 0-0.019% of total B cells from 4 hours to 7 days post-infection) but readily detected (median 0.81%) by day 14 (**Fig. 1b, Extended Data Fig. 1b**). While organoids from all donors (n=40) could be infected, their frequencies varied substantially, demonstrating host heterogeneity in the propensity for EBV infection or expansion of infected cells (**Fig. 1c**).

**Figure 1:**
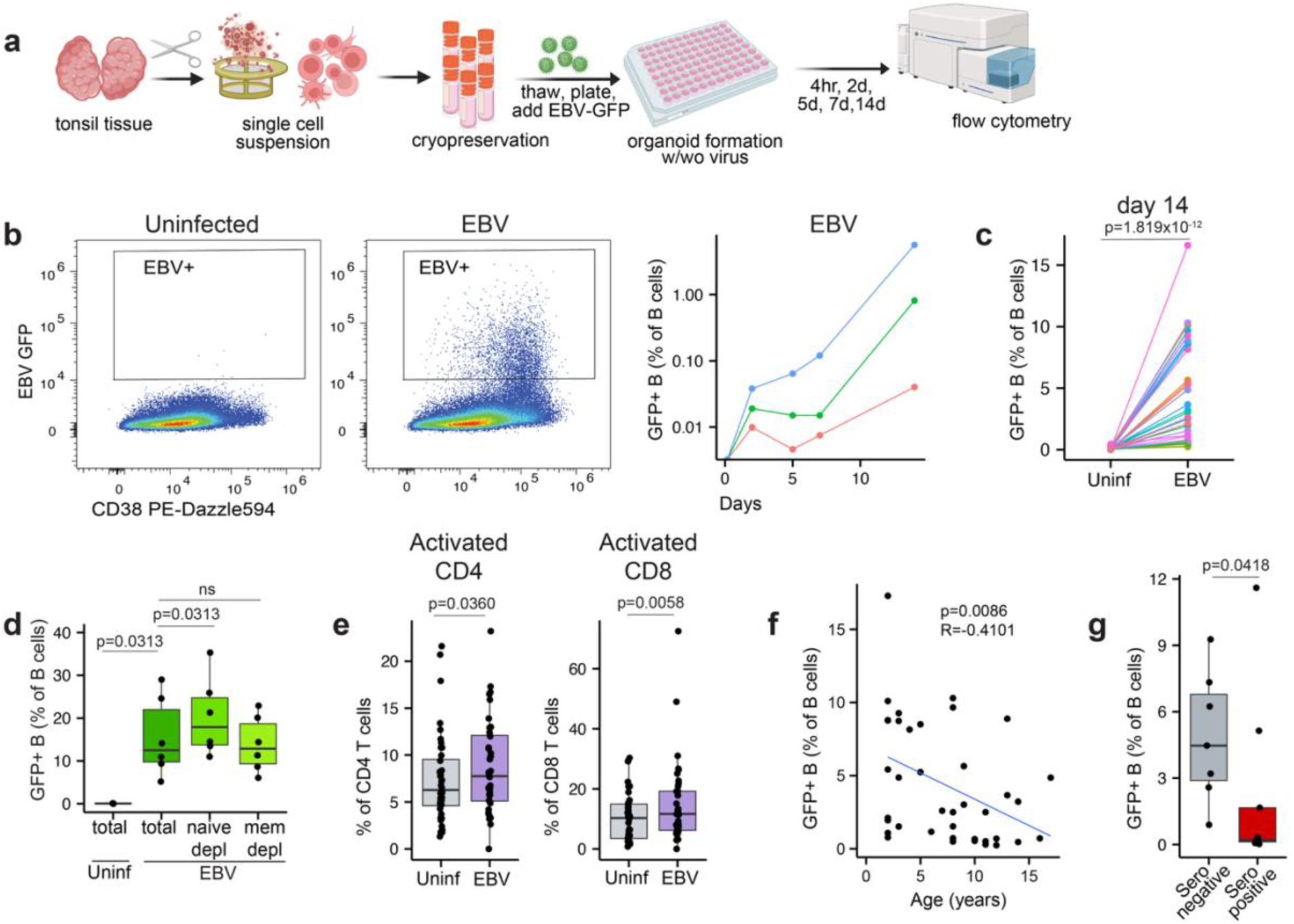
Assessment of EBV^+^ B cells in the tonsil organoids. (a) Workflow. Immune organoids were prepared from tonsils and infected with 0.000005 or 0.00005 or 0.0005 MOI of GFP-expressing EBV (Akata strain). EBV-infected and uninfected organoids were harvested 4 hours, 2 days, 5 days, 7 days, and 14 days post infection and evaluated by flow cytometry. Image created in BioRender. (b) Representative flow cytometry plots for EBV^+^ B cells on day 14 and kinetics of EBV^+^ B cells in response to 0.0005 MOI of GFP-expressing EBV (n=3 per group). (c) EBV^+^ B cell frequencies in tonsil organoids on day 14 post infection (n=40 per group). (d) EBV^+^ B cell frequencies on day 14 in uninfected and EBV-infected organoids in the presence or absence of naive (CD38^-^CD27^-^IgD^+^) or memory (CD38^-^CD27^+^IgD^-^ and CD38^-^CD27^-^IgD^-^) B cells (n=6 per group). (e) Activated (CD38^+^HLA^-^DR^+^) CD4 and CD8 T cell frequencies in the uninfected and EBV-infected organoids on day 14 (n=40 per group). (f) Correlation between tonsil donor age and EBV^+^ B cell frequencies (n=40 per group). (g) Frequency of EBV^+^ B cells on day 14 in EBV-infected tonsil organoids derived from EBV seronegative and seropositive individuals (n=16; 7 seronegative, 9 seropositive). Mann Whitney U tests were used to calculate p values between groups. Spearman’s rank test was used for correlation analysis. Boxplots indicate the median value, with hinges denoting the first and third quartiles and whiskers denoting the highest and lowest value within 1.5 times the interquartile range of the hinges. MOI, multiplicity of infection; ns, not significant (p ≥ 0.05).

EBV can immortalize a range of B cells, including naive, unswitched, and switched memory B cells^40–42^. Naive B cells are thought to be a major EBV target *in vivo*, whereas resting memory B cells are the persistent reservoir^6,8^. However, it has been difficult to establish which subsets are initially infected and how they differentiate, such as by GC or extrafollicular pathways, prior to entering the memory pool^42–44^. To evaluate which B cells are permissive to EBV infection in the tonsil organoid microenvironment, we depleted either memory (CD27^+^CD38^-^IgD^-^ and CD27^-^CD38^-^IgD^-^) or naive (CD27^-^CD38^-^IgD^+^) B cells on day 0, infected with EBV, and later quantified GFP^+^ B cells compared to non-depleted control organoids. Naive B cell depletion led to a significant increase in EBV^+^ B cells, whereas depletion of memory B cells had no impact (**Fig. 1d**). These results indicate that while both naive and memory B cells are permissive to EBV infection, memory B cell infection and/or outgrowth may be greater than that of naive B cells in a secondary lymphoid microenvironment.

Since T cell responses can restrain EBV^+^ B cell outgrowth, we examined T cell activation (based on CD38 and HLA-DR co-expression) in infected organoids. Total CD4 and CD8 T cell frequencies were unchanged by EBV infection (**Extended Data Fig. 1c**) but activated phenotypes were modestly but significantly higher in EBV-infected organoids (**Fig. 1e**). EBV reactivation is restrained by cellular responses *in vivo*^25,45,46^. We therefore tested whether tonsil donor age and/or EBV seropositivity (as indicated by antibodies against viral capsid antigen (VCA) and Epstein-Barr nuclear antigen (EBNA) in matched serum) were associated with organoid EBV^+^ B cell proportions. Indeed, organoids from older donors (**Fig. 1f**; p=0.0086) and from EBV seropositive donors had fewer EBV^+^ B cells (**Fig. 1g**; p=0.0418), suggesting that organoids can capture recall responses that limit B cell infection and/or expansion.

#### Tonsil organoid EBV^+^ B cells acquire a GC-like phenotype

We next characterized the phenotypes of EBV^+^ B cells over the first two weeks of culture. Based on CD38, CD27, and IgD patterns, which are commonly used to assign B cell states, most EBV^+^ B cells acquired activated (preGC) or GC-like phenotypes by day 14 (**Fig. 2a**). A few EBV-infected B cells developed plasmablast-like phenotypes. These profiles were specific to tonsil organoids, as parallel organoids experiments using peripheral blood mononuclear cells (PBMCs) or splenocytes as source material showed either inefficient EBV infection (for PBMCs) or fewer pre-GC B cell phenotypes (for splenocytes) compared to tonsils (**Extended Data Fig. 2a**). In tonsil organoids, we also evaluated the relative contribution of naive vs. memory B cells to the EBV^+^ B cell differentiation phenotypes by individually depleting these subsets prior to establishing organoids. EBV^+^ GC-like and plasmablast B cells were elevated when naive B cells were depleted, while infected B cells with a preGC phenotype were unaltered by depletion (**Fig. 2b**). By contrast, organoids established from memory B cell-depleted tonsils exhibited slightly lower levels of EBV^+^ plasmablasts, but pre-GC and GC populations were unaltered. These results suggest that memory B cell infection may be an important contributor to the EBV^+^ B cell GC and plasmablast tonsil reservoir.

**Figure 2:**
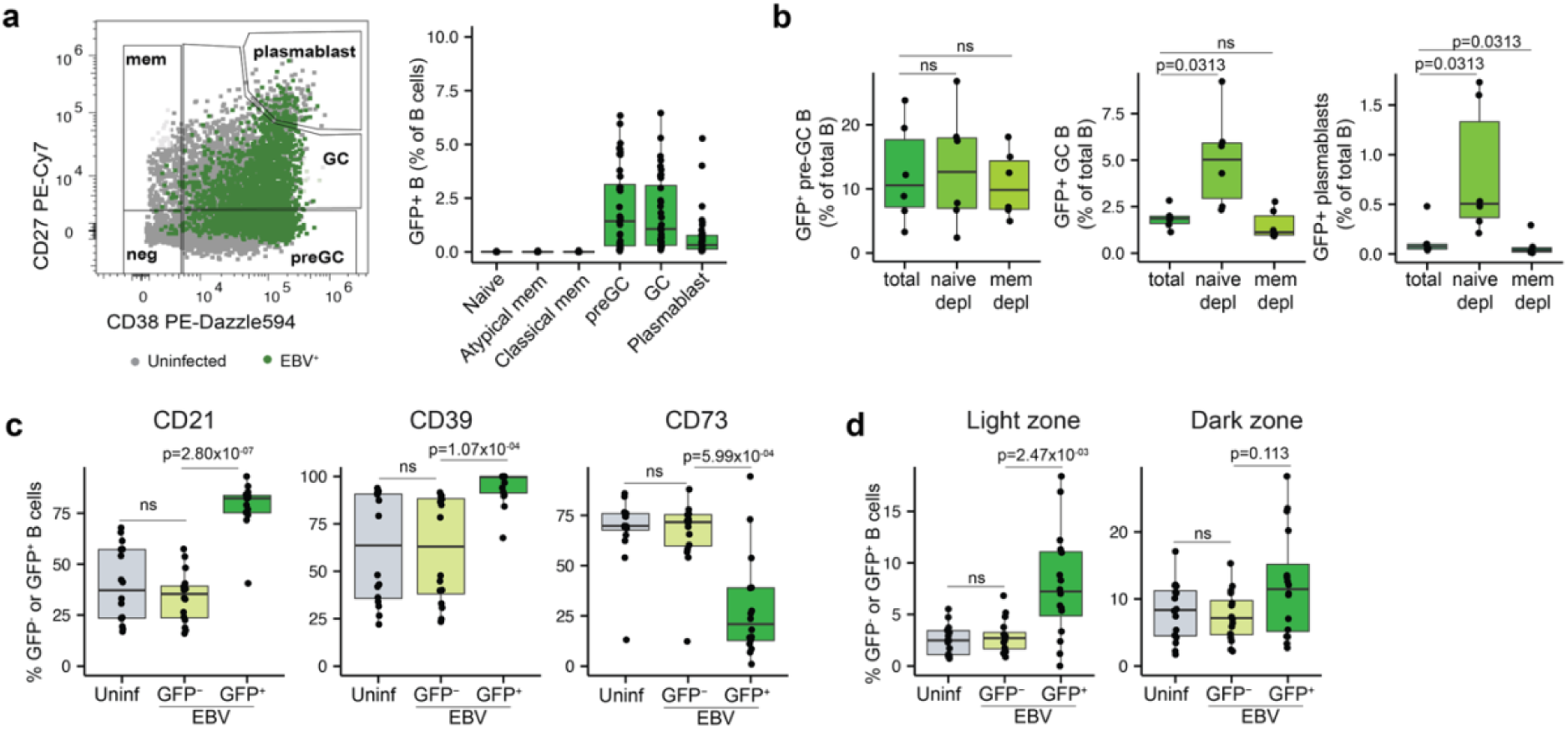
Phenotypes of EBV^+^ B cells in tonsil organoids. (a) Representative flow cytometry plot showing EBV^+^ (green) and uninfected (grey) B cells and their frequencies (right) in tonsil organoids on day 14 post infection (n=40 per group). (b) Phenotypes (on day 14) of EBV^+^ B cells when pre-existing naive or memory B cells were depleted at the initiation of organoid culture (n=6 per group). Expression of (c) CD21, CD39, CD73 and (d) light (CD83^+^) and dark (CXCR4^+^) zone markers in uninfected organoids and GFP^-^ and GFP^+^ B cells from EBV-infected organoids (n=16 per group). Mann Whitney U tests were used to calculate p values between groups (b) and where appropriate, multiple hypothesis correction using the Benjamini & Hochberg method was performed (c, d). Boxplots indicate the median value, with hinges denoting the first and third quartiles and whiskers denoting the highest and lowest value within 1.5 times the interquartile range of the hinges. ns, not significant (p ≥ 0.05).

We further characterized other cell surface markers on EBV^+^ B cells that are conventionally used to define lymphoid tissue B cell subsets. Relative to uninfected cells, EBV^+^ B cells were more likely to be CD21^+^, a gp350 attachment receptor for EBV^47^, and CD39^+^, an enzyme that hydrolyzes extracellular ATP and ADP^48^ (**Fig. 2c**). In contrast, CD73, an enzyme responsible for producing immunosuppressive adenosine^49^, was significantly reduced on EBV^+^ cells. Taken together, these results suggest EBV^+^ B cells may modulate extracellular ATP to AMP, potentially for immunomodulation.

The GC is compartmentalized into light and dark zones, where antigen-activated B cells affinity mature, proliferate and mutate their B cell receptors (BCRs)^43^. Although the overall proportions of light and dark zone B cells were unaffected by EBV infection (**Extended Data Fig. 2b**), EBV^+^ B cells were enriched for expression of the light zone marker CD83 (p=0.00247) but not for dark zone marker CXCR4 (p=0.113) (**Fig. 2d**). Together, these phenotypic differences suggest that early following infection, EBV^+^ B cells have distinct activation, proliferation, and immunosuppressive properties^50^ compared to their uninfected counterparts.

#### Primary EBV infection elicits unique and diverse transcriptional tonsillar B cell states

Although several prospective studies have followed EBV-naive individuals through initial infection^51,52^, primary responses to EBV remain incompletely defined, in part because early infection is often asymptomatic, particularly in young individuals. In symptomatic IM cases, sampling typically happens 4-8 weeks post-infection^53,54^. Therefore, we leveraged the tonsil organoid model to perform a detailed longitudinal analysis of the early events of primary EBV infection in seronegative donors (**Extended Data Fig. 3a**). At single cell resolution, we measured gene expression, B and T cell receptor repertoires, and selected protein expression over the first three weeks of infection in organoids derived from four EBV-naive donors. Aggregation of 89,444 cells from all donors, infection conditions, and time points revealed 18 main cell clusters **(Extended Data Fig. 3b,c; Extended Data Table 2**), which were then sub-divided by cell identity for additional analyses.

We first evaluated the B cell transcriptional landscape following EBV infection. Sub-clustering defined 21 B cell clusters (**Extended Data Table 3)** and *GFP* mRNA was detected in several of them, indicating EBV^+^ B cells were present across numerous cell states (**Fig 3a, Extended Data Fig. 3d**). Stratification by organoid infection and time revealed a dramatic, EBV-induced transcriptional change in B cells on days 14 and 21 post-infection (**Fig 3b**). Further analysis of the top differentially expressed genes (DEGs) at later time points showed remarkable transcriptional diversity in EBV^+^ B cells and almost no transcriptional overlap with uninfected cells (**Fig. 3c,d**; EBV-associated clusters bolded). At a broad level, the 11 EBV-associated clusters spanned GC, plasmablast, and proliferating phenotypes (**Fig. 3c,d**, **Extended Data Fig. 3e**) and the proportions of these clusters increased over time (**Fig. 3e**, **Extended Data Fig. 3f**).

**Figure 3:**
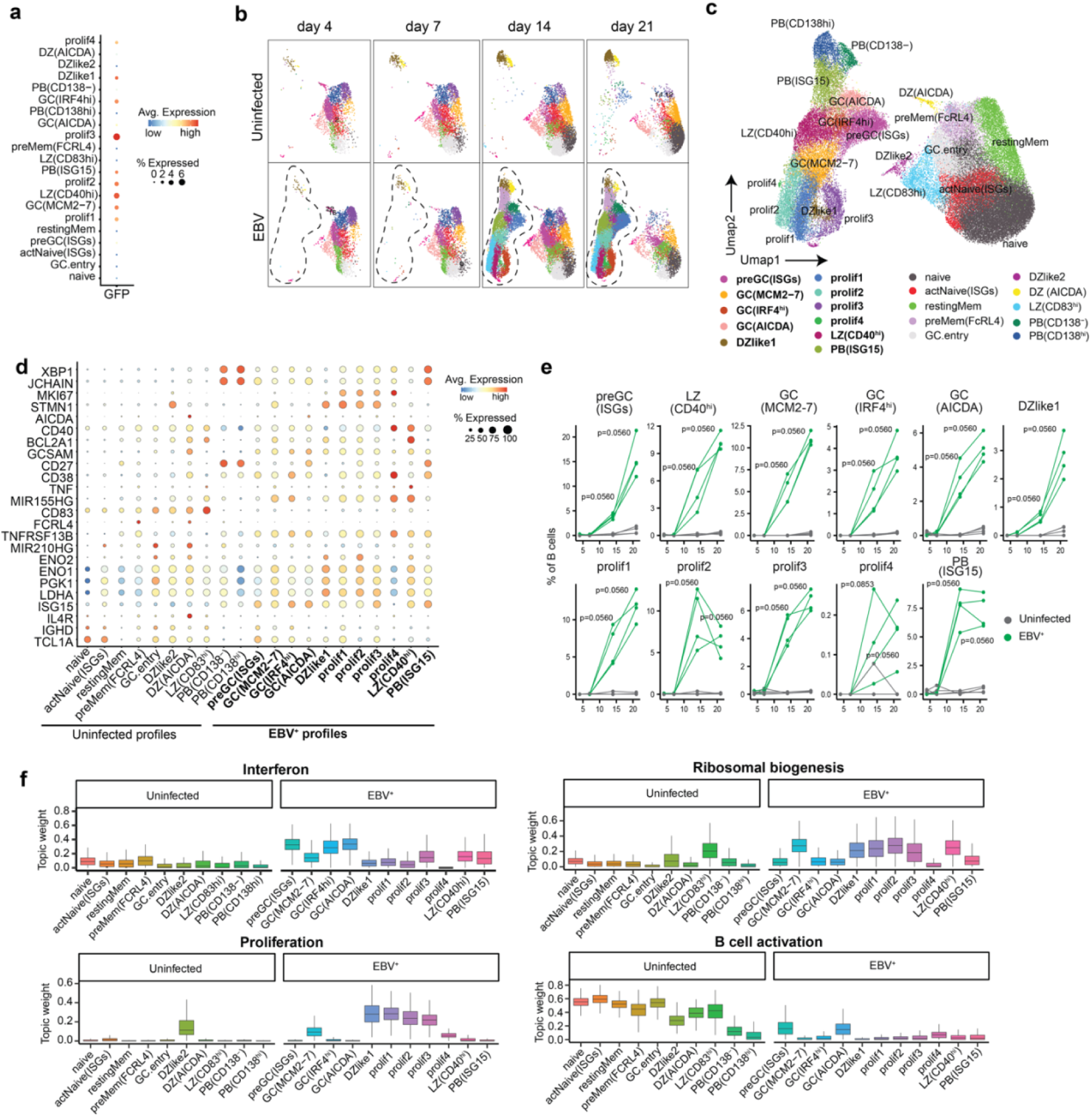
Transcriptional landscape of EBV^+^ B cells in primary infection. (a) Detection of GFP transcripts in B cell clusters from EBV-infected organoids. (b) UMAP stratification by infection condition and time. Populations enriched following EBV infection are highlighted. (c) UMAP of B cells (all donors, timepoints and stimuli combined, 63,263 B cells, n=4 seronegative donors, 1 experiment). B cell clusters associated with EBV infection in organoids are bolded. (d) Differentially expressed genes used to define B cell cluster identities. (e) Frequencies of B cell clusters in uninfected (grey) and EBV^+^ (green) organoids (n=4 per group). (f) Topic weights (mean gene expression program) across B cell clusters in EBV-infected organoids. Boxplots indicate the median value, with hinges denoting the first and third quartiles and whiskers denoting the highest and lowest value within 1.5 times the interquartile range of the hinges. Mann Whitney U tests followed by multiple hypothesis correction using the Benjamini & Hochberg method were used to calculate p values.

We noted several unique features of the EBV^+^ populations. Even though many GC-like profiles were present in the EBV^+^ B cell pool, most had low expression of the master transcription factor *BCL6*, as has been noted in previous studies^55,56^. Some of the GC-like, EBV^+^ B cells were primarily defined by conventional GC gene expression features, such as *MCM2-*7 (involved in DNA replication) and *AICDA* (encoding AID, the enzyme that facilitates somatic hypermutation and class switching). However, the top DEGs for several EBV^+^ GC populations were dominated by interferon (IFN) response signaling, such as IFN-stimulated genes *(ISGs)* and IFN regulatory factors (e.g. *IRF4*) (**Fig. 3d**), demonstrating they were responding to surrounding antiviral signals. Within plasmablasts, an EBV^+^ subset was also strongly defined by *ISG* expression signatures (**Fig. 3c-e, Extended Data Fig. 3e**). Aside from these populations, a substantial proportion of EBV^+^ B cells expressed *MKI67 and STMN1 (*Prolif1-4), indicating a large fraction were proliferating; this proliferation was largely restricted to infected cells. We did not identify any naive or memory B cell subsets in the EBV^+^ pool (**Fig. 3c-e**, **Extended Data Fig. 3e,f)**, suggesting EBV drove differentiation away from a naive state but not yet to a memory state by day 21 post-infection. Overall, the heterogeneity of GC-like B cell subsets highlights the diversity of EBV-induced B cell transcriptional reprograming within the secondary lymphoid nice.

By day 14, organoid EBV^+^ B cells exhibited diverse transcriptional states, indicating that the virus exploits multiple functional niches to drive proliferation and survival while bypassing canonical GC checkpoints, thereby supporting viral persistence. We performed topic analysis^57,58^ to summarize key EBV-driven transcriptional remodeling. We identified 15 topics across infected vs. uninfected B cell populations (**Extended Data Fig. 3g**, **Extended Data Table 4**). Classification based on top 50 DEGs highlighted IFN response (K6), ribosomal biogenesis (K11), and proliferation (K15) topics in EBV^+^ B cells (**Fig. 3f**). The IFN topic was most evident in GC-like EBV^+^ B cells but also gradually increased in bystander B cells at later time points (**Extended Data Fig. 3h, Extended Data Table 5**). Ribosomal biogenesis was particularly increased in EBV^+^ proliferating populations (**Extended Data Fig. 3h, Extended Data Table 5**), indicative of rapid proliferation. Our data reveal that EBV selectively promotes growth in B cells with low expression of the canonical GC regulator *BCL6* in the tonsil microenvironment. This decoupling of proliferation from normal GC checkpoints may point to a mechanism by which EBV expands its reservoir while evading standard GC-mediated regulation.

#### A small subset of EBV^+^ B cells exhibit conventional GC B cell activity

It remains controversial whether EBV^+^ B cells participate in GC reactions, in which B cells proliferate, undergo SHM and CSR, and differentiate into memory or plasmablast phenotypes. While EBV^+^ B cells have been identified in tonsillar GCs^15,18^, they do not show evidence of SHM activity, at least in IM patient tonsils^20^. Yet, IM patients have abundant class-switched memory B cells with evidence of SHM^19^, suggesting either GC transit, direct memory B cell infection^41^, or an EBV-driven mechanism different from conventional B cell differentiation. Recent work indicates EBV^+^ B cells may phenocopy GC dynamics, although they are transcriptionally distinct from conventional tonsillar GC B cells^59^. To reconcile these findings and address the plausibility of EBV^+^ B cells participating in an active GC response, we performed B cell receptor (BCR) and isotype analysis from tonsil organoids. On day 14 post-infection, ∼40% of EBV^+^ B cells had class switched, as compared with 10-15% of uninfected B cells from the same organoids and uninfected control organoids (**Fig. 4a,b)**. We hypothesized that either class-switched B cells had a higher propensity for infection, or that EBV infection enhanced class switching through an unknown mechanism without increasing SHM. To differentiate between these two possibilities, we looked for evidence of SHM in switched and unswitched B cell populations, since on average *ex vivo* class-switched B cells would have higher SHM rates than unswitched cells. Analysis of SHM rates by isotype revealed that on average, EBV^+^ cells had similar or even lower SHM rates compared to uninfected B cells of the same isotype (**Fig. 4c, Extended Data Fig. 4a**). The only exception to this was the IgM population, where EBV^+^ cells had slightly higher SHM, though rates were, as expected, very low across all conditions (**Extended Data Fig. 4a)**. These findings are consistent with EBV-induced class switching of naive B cells without, for the most part, induction of SHM in the tonsil organoid microenvironment.

**Figure 4:**
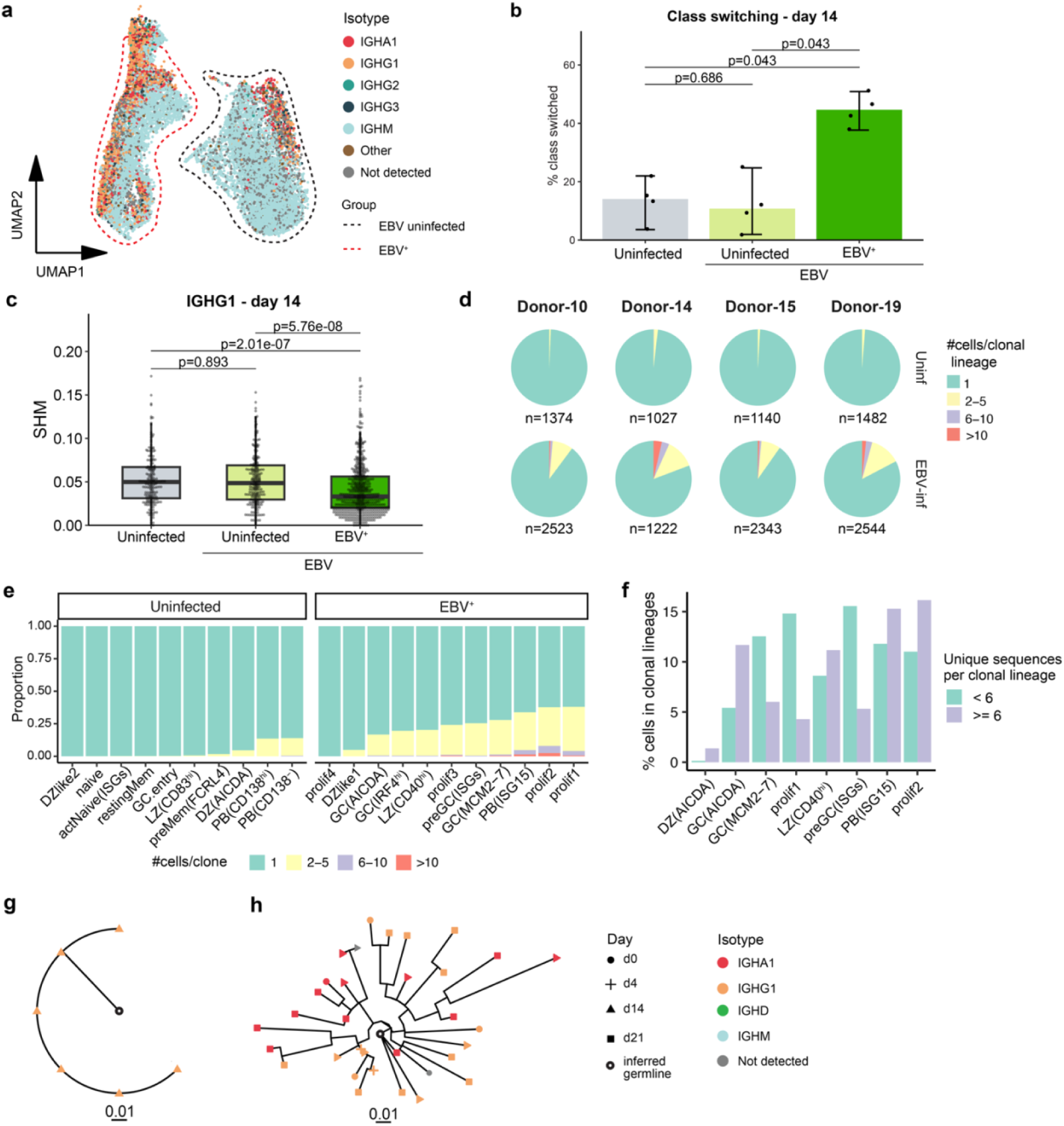
Clonality and isotype usage of EBV^+^ tonsil organoid B cells. (a) Antibody isotype usage in EBV^+^ and uninfected B cells from EBV-infected organoids in four donors on day 14. Isotype usage was mapped onto the B cell UMAP generated in Fig 3. (b) Frequency of class-switched isotypes in uninfected, EBV-uninfected and EBV^+^ B cells on day 14. (c) SHM frequency (95% CI) among IGHG1 B cells in uninfected, EBV-uninfected and EBV^+^ B cells on day 14. (d) BCR repertoire diversity in EBV-infected and uninfected organoids from EBV-seronegative donors on day 14. Clonal lineages were defined, then the number of cells per clonal lineage were plotted as a proportion of total B cells. (e) Transcriptional profiles associated with clonal expansion. Most expanded clones are in EBV-associated B cell transcriptional profiles. Data from day 14 post-infection. (f) Size distribution of clonal lineages containing at least one EBV^+^ B cell. Clonal lineages with more diverse BCR representation (purple) were found primarily in GC-associated cluster phenotypes. B cell cluster frequencies present in proliferating and GC-like clonal lineages from ex vivo cells and EBV-infected organoids. Lineages (n=160) with more than 15 cells are shown. Examples of clonal lineages in EBV^+^ B cells from (g) a simple clonal expansion following infection, and (h) an elaborate GC-like clonal lineage with greater BCR diversification, SHM, and class switching. 0.01 indicates the branch length representing the number of mutations per site between each node in the tree. Boxplots indicate the median value, with hinges denoting the first and third quartiles and whiskers denoting the highest and lowest value within 1.5 times the interquartile range of the hinges. Mann-Whitney U tests (two-sided) followed by Benjamini & Hochberg method were used to calculate significance values (*p<0.05, **p<0.01, ***p<0.001, ****p<0.0001). SHM, somatic hypermutation; BCR, B cell receptor; CI, confidence interval.

We also analyzed BCR repertoire diversity in EBV^+^ vs. uninfected organoids from the EBV seronegative donors on day 14 post-infection. Consistent with increased frequency of EBV^+^ GC and proliferating cell profiles, EBV-infected organoids had more B cell clonal expansions and clonal lineages (**Fig. 4d**). Clonal expansions were enriched within EBV^+^ pre-GC, GC, and proliferating clusters (**Fig. 4e**), suggesting that EBV drove proliferation within multiple infected B cell states, including prior to GC entry. The largest and most diverse clonal lineages were observed in EBV^+^ GC-like B cells (**Fig. 4f**). Although 99% (3408 out of 3437) of EBV^+^ clonal lineages with at least six B cell members were simple expansions with limited SHM (**Fig. 4g)**, we found several examples (n=29 lineages, 0.84%) of elaborate clonal lineages with SHM (**Fig. 4h, Extended Data Fig. 4b**) that were unique to the EBV^+^ B cell population. Cells from these larger lineages were detected across numerous time points and showed evidence of intra-lineage class switching, consistent with GC functions **(Fig. 4h)**. Based on these findings, we conclude that EBV infection supports isotype class switching and most EBV^+^ B cells undergo small, simple clonal expansions without BCR mutation, but a small subset of infected B cells do undergo SHM in the tonsil organoid microenvironment.

#### CD4 T cells limit EBV^+^ B cells in tonsil organoids

CD4 and CD8 T cells contribute to EBV control^11,22,60,61^, though little is known about initial T cell responses to EBV infection, particularly in the tonsil. A major benefit of the tonsil organoid model is the ability to holistically consider both B and T cells in the same culture environment. Thus, we analyzed CD4 and CD8 T cell transcriptional profiles in the same EBV seronegative donor-derived organoids (**Extended Data Fig. 3a)**. Aggregation and analysis of *CD3^+^* cells (18,707) from all donors, time points, and infection conditions identified 15 T cell clusters (**Fig. 5a, Extended Data Fig. 5a,b; Extended Data Table 6**). Overall, the populations identified were in line with our previous studies of *ex vivo* and organoid-cultured tonsil T cells^38^. In contrast to our B cell analyses, we did not detect any T cell clusters uniquely associated with or induced by EBV (**Fig. 5b**). Rather, the proportion of T cells occupying different clusters shifted over time. By day 14, EBV-infected organoids showed a trend (though not statistically significant due to small sample size) for increased proportions of activated CD4 T cells and germinal center T follicular helper (GC-TFH) cells with *ISG* expression profiles, coinciding with EBV^+^ B cell outgrowth (**Fig. 5c**). Minor increases in these populations, in addition to a proliferating T cell population, started on day 4 and peaked on day 14 post-infection (**Fig. 5c**). Other T cell populations, including memory CD8 T cell subsets conventionally associated with EBV infection in peripheral blood studies, were largely unchanged following tonsil organoid infection (**Extended Data Fig. 5c**). This disparity may be due to the exceptionally low dose used to infect the organoids and the kinetics of a naive EBV response. As a complementary analysis, secreted cytokines were quantified from organoid supernatants. Among the 36 cytokines quantified, IFN2α, MIP-1α, RANTES, TNF-β were significantly elevated in EBV-infected organoids (**Fig. 5d**). IL-10 is an important cytokine in EBV infection, and the virus encodes a homolog to modulate IL-10 signaling; we saw a small, but not significant increase in IL-10 levels following EBV infection. Overall, both proinflammatory and anti-inflammatory cytokines were induced in organoids in response to EBV.

**Figure 5:**
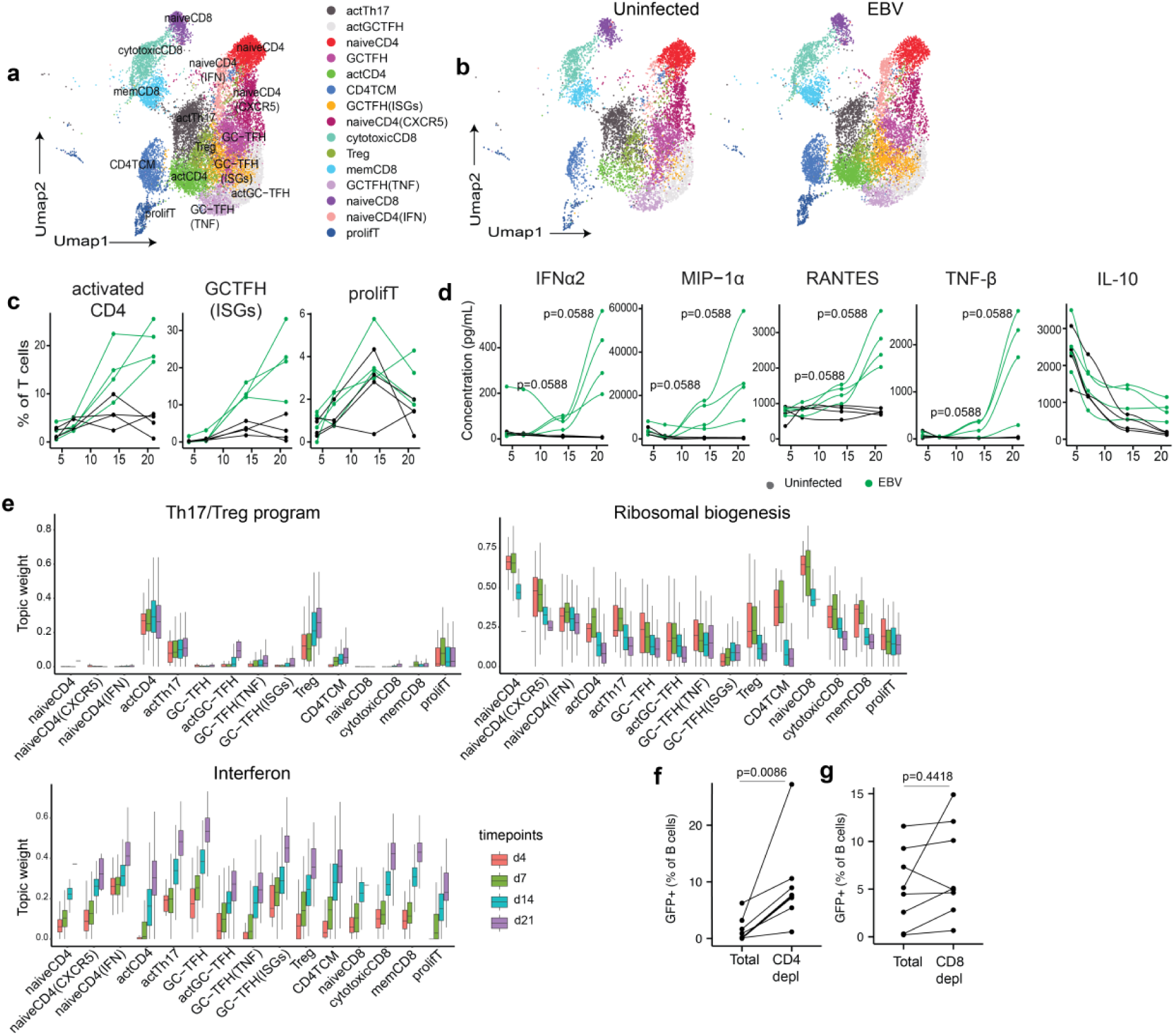
T cell responses to EBV in tonsil organoids. (a) Aggregated UMAP representing T cell subsets from EBV seronegative donors (all donors, timepoints and stimulation aggregated;18,707 cells, n=4, 1 experiment). (b) Distribution of T cell transcriptional profiles in uninfected and EBV-infected organoids. (c) Frequencies of activation-associated CD4 T cell clusters over time in EBV-infected (green) and control (grey) organoids (n=4 per group). No cluster frequencies reached statistical significance between uninfected and EBV-infected organoids. (d) Secreted cytokines in uninfected (grey) vs. EBV-infected (green) organoids (n=4 per group). (e) Topic weights (mean gene expression program) in CD4 and CD8 T cell subsets in EBV-infected organoids. EBV^+^ B cell frequencies in the organoids on day 14 in the presence or absence of (f) CD4 T cells (n=8 per group) and (g) CD8 T cells (n=8 per group). Boxplots indicate the median value, with hinges denoting the first and third quartiles and whiskers denoting the highest and lowest value within 1.5 times the interquartile range of the hinges. Mann Whitney U tests followed by multiple hypothesis correction using the Benjamini & Hochberg method were used to calculate p values (c-e). Mann Whitney U tests were used to calculate p values between groups (f,g).

Topic modeling was performed to further characterize changes in T cell states following EBV infection. Eight topics were identified across the T cell dataset (**Extended Data Fig. 5d**, **Extended Data Table 7**). Annotated topics of interest included a Th17/Treg program (k1), protein synthesis/ribosomal biogenesis (k2), cytotoxicity (k3), proliferation (k4), interferon response (k6), and hypoxia/apoptosis (k7). Most topics corresponded well with our prior T cell annotations based on top DEGs except for the Th17/Treg topic (k1), which was significantly enriched in activated CD4, Th17, and Treg clusters in infected compared to uninfected organoids (**Fig. 5e, Extended Data Table 8**). CD4 T cell polyfunctionality has been described in response to EBV; it is plausible that these CD4 T cell subsets exhibit concurrent pro- and anti-inflammatory features. We observed significant induction of the IFN-related topic across all T cell subsets in infected organoids (**Fig. 5e, Extended Data Table 8**), suggesting a strong and early antiviral response. In contrast to B cells, the ribosomal biogenesis topic was significantly downregulated across many CD4 and CD8 T cell populations following infection (**Fig. 5e**), perhaps due to IFN-induced antiviral mechanisms. We assessed whether EBV infection perturbed T cell repertoire diversity and found there were no substantial changes in CD4 or CD8 T cell clonality (**Extended Data Fig. 5e**). T cells likely have distinct functions in acute vs. chronic phases of EBV infection and it is possible that clonal expansions would be observed beyond day 21. Finally, we tested the ability of CD4 vs. CD8 T cells to control EBV infection through depletion studies in a separate cohort of tonsillectomy samples with mixed serostatus for prior EBV exposure. CD4 T cell depletion led to a significant increase in EBV^+^ B cells by day 14 post infection (**Fig. 5f**). In contrast, CD8 T cell depletion did not substantially change EBV^+^ B cell proportions in tonsil organoids (**Fig. 5g**). Overall, these results highlight a major role for CD4 T cells in controlling EBV^+^ B cell outgrowth in the tonsil organoid microenvironment.

### Discussion

EBV is a strictly human pathogen and while major progress has been made, aspects of acute and chronic infection remain difficult to model in animal models. To complement these efforts, human tonsil organoids were used to investigate primary EBV infection. EBV^+^ B cells were phenotypically and transcriptionally distinct from uninfected cells, including numerous and diverse cell states not previously reported. EBV^+^ B cells exhibited a greater propensity to class switch but had comparatively low SHM rates. Although many EBV^+^ cells showed signs of proliferation and underwent small clonal expansions, only a tiny fraction of B cell clones showed SHM patterns consistent with genuine GC participation. Along with B cell studies, we defined features of the early T cell response to EBV infection. Based on these findings, we present a working model of the dynamics of initial EBV infection in the tonsil (**Fig. 6**) and our data suggest tonsil organoids are a physiologically relevant model for studying EBV biology in a relevant human tissue context.

**Figure 6:**
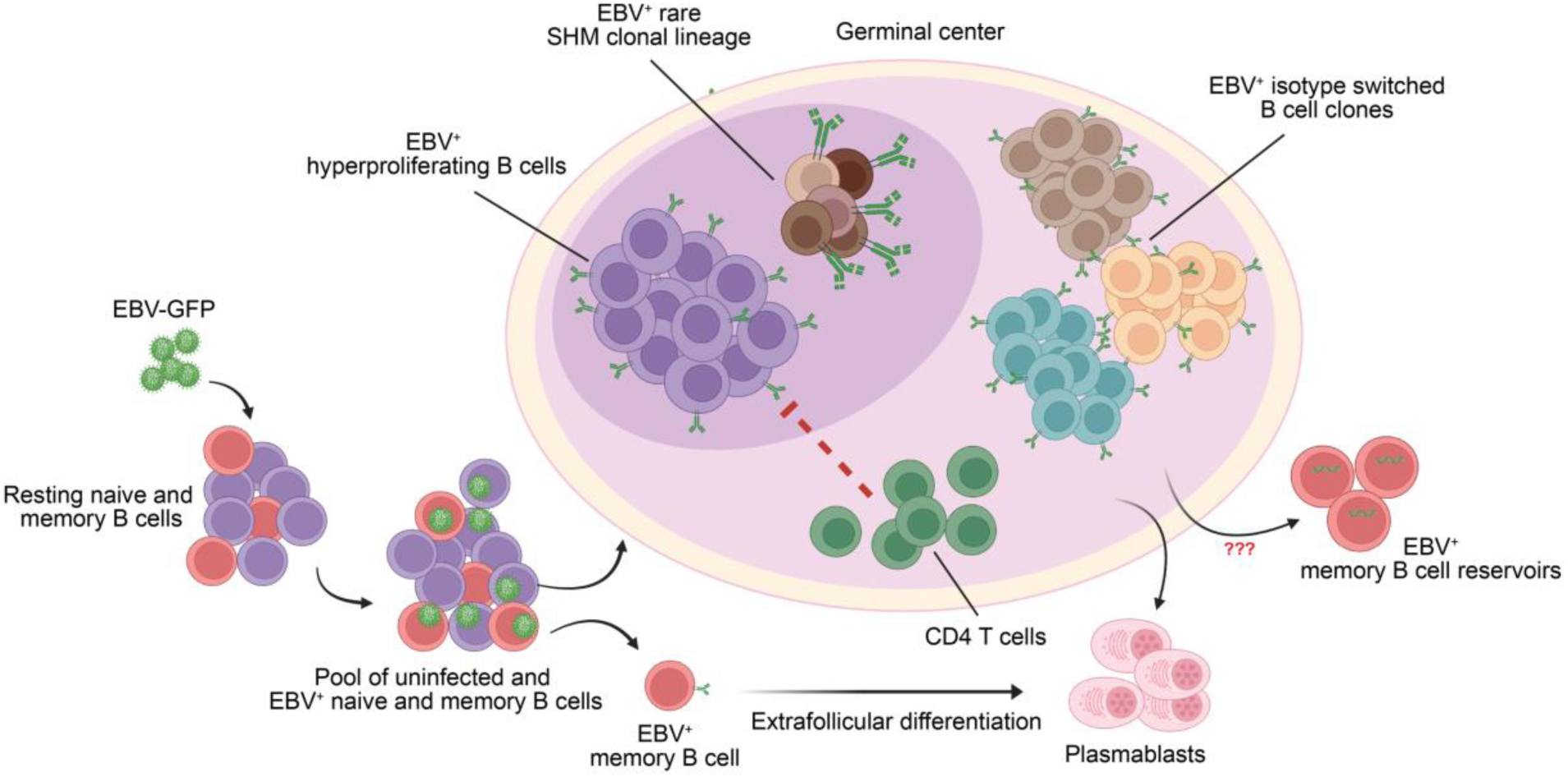
A working model of primary EBV infection in human tonsil organoids. Upon infection, B cells in tonsil organoids acquire GC-like phenotypes. Most infected cells class switch and undergo modest proliferation without BCR mutation, leading to many small clonal expansions. A tiny fraction of EBV^+^ GC B cells engage in somatic hypermutation and develop elaborate clonal lineages. Both populations can differentiate into EBV^+^ plasmablasts although differentiation into long-term resting memory B cells remains unknown. CD4 T cells inhibit EBV^+^ B cell outgrowth. Image created in BioRender.

The GC model of EBV infection proposes that *in vivo*, EBV^+^ naive B cells transit through the GC to establish latency in the memory B cell pool. In support of this hypothesis, EBV^+^ B cells have been identified in the GC and within interfollicular regions of latently infected tonsils^17^, where they are posited to undergo varying degrees of proliferation balanced by cell death. More recently, they have also been identified in the marginal zone^62^. However, whether the EBV^+^ B cells participate in a *bona fide* GC reaction has remained uncertain^18,20^, particularly in the setting of acute infection. We now provide experimental evidence that EBV^+^ B cells can indeed participate in a genuine GC reaction in a secondary lymphoid model. This was underscored by the identification of a small subset of EBV^+^ B cells with extensive SHM and clonal expansion, highly suggestive of transit through a genuine GC. Such sporadic participation of the EBV^+^ B cells in a true GC reaction offers experimental support for the EBV GC model. We also found that EBV could directly infect either naive or memory B cells, with memory B cells showing enhanced infectivity in the absence of naive B cells, suggesting the propensity for naive cells to be the major initial target of infection may be a matter of abundance. These findings raise the possibility that in addition to transiting through the GC, EBV may also access the memory compartment through direct memory B cell infection.

At the transcriptional level, EBV^+^ B cells were transcriptionally heterogeneous and distinct from uninfected cells. This diversity likely reflects viral manipulation of GC-associated programs to accommodate immune pressure while promoting infected cell survival. Notably, EBV-induced GC-like B cell subsets expressed low levels of *BCL6*, the master regulator of GC B cells, consistent with prior reports demonstrating *EBNA3C* and EBV miRNA-mediated suppression of BCL6^55,56,63,64^. Within the GC, the light zone mediates selection and affinity maturation. Flow cytometric analysis revealed that EBV^+^ GC B cells were enriched for a light zone phenotype, while dark zone frequencies were comparable between EBV^+^ and uninfected populations. EBV latent membrane proteins LMP1 and LMP2A mimic CD40 and BCR signaling, respectively, potentially enabling EBV^+^ B cells to persist in a light zone-like state independent of high-affinity antigen^65–67^. Accordingly, enrichment of light zone-like cells among EBV^+^ B cells suggests a preferential survival strategy, potentially involving enhancer-mediated upregulation of *BCL2A1*^68^. In contrast, the unchanged frequency of dark zone B cells supports EBV-mediated subversion of the conventional affinity maturation cycle, consistent with our BCR sequencing data showing CSR in the absence of SHM for most infected B cells.

Although EBV can upregulate AID, the enzyme that mediates SHM and CSR^69,70^, these processes are mechanistically and spatially separable. *AICDA* mRNA was detected in one EBV^+^ GC-like subset. Yet, most EBV^+^ B cells class switched without undergoing SHM. This is consistent with limited data from IM patients, which found that EBV^+^ B cells have higher levels of CSR than uninfected B cells^71^. Notably, CSR can be induced by strong CD40 signaling or by its EBV mimic LMP1, which can promote extrafollicular CSR independent of cognate T cell help^72–74^. Such uncoupling of CSR from SHM may favor EBV persistence by limiting potentially deleterious mutagenesis of the double stranded DNA viral genome, which could otherwise be a consequence of extensive SHM. Along with CD40 signaling, cytokines including IL-10, IL-4, and TGF-β are known regulators of CSR^76^. In addition to inducing host IL-10 expression, the EBV encoded IL-10 homolog BCRF1^77^ may promote CSR, perhaps through leaky lytic gene expression. The relative contributions of host-derived cytokines and viral cytokine homologs to EBV-driven CSR remain to be determined and warrant further investigation.

While EBV^+^ B cell differentiation into memory cells has been observed *in vitro*^21^, we did not identify EBV^+^ memory B cells in the organoids. It is possible that the 3-week culture period was not sufficient to establish a pool of EBV-infected memory B cells. Alternatively, EBV^+^ memory B cells may have silenced all EBV gene expression and become indistinguishable from conventional, uninfected memory B cells in the organoids. Future studies will delineate the temporal dynamics of EBV gene expression in the tonsil organoid system to associate viral gene expression patterns with host B cell transcriptional states during infection.

CD4 and CD8 T cells exert anti-EBV responses to both latent and lytic antigens^25^. CD4 T cells specific for peptides from EBV latency proteins exhibit Th1-like profiles but without overt expansion of the CD4 T cell compartment^78^. EBV-specific CD8 T cells often target lytic antigens in early IM^22,79^ and in IM patients, CD8 T cells specific for EBV lytic and latent peptides can represent 1-40% and 0.1-5% of total circulating CD8 T cells, respectively^24,80,81^. Activated CD4 and CD8 T cell frequencies each increased in response to EBV in the tonsil organoids, though we observed relatively stable CD4 and CD8 T cell repertoire diversity despite CD4 T cell transcriptional activation and T cell-derived cytokine production. The limited CD8 T cell activation may reflect the extremely low viral dose used to infect the organoids, low lytic replication, as well as the young age of the EBV-seronegative tonsil donors (<13 years), in whom primary EBV infection is often clinically mild or asymptomatic. Nonetheless, the tonsil organoid model offers further opportunities to study anti-EBV T cell responses in primary vs. secondary infection, including mechanisms of EBV immune evasion.

In children and adults with primary EBV infection, granzyme B and perforin-secreting CD4 T cells have been identified in tonsils and peripheral blood, respectively^46,82^ and the cytotoxic profile of Th1-like CD4 cells is retained in the tonsil throughout the lifetime^83^. LMP1 signaling induces potent T cell responses, including cytotoxic CD4 T cells^60,74^. Polyfunctional T cells may help to control chronic infection^84^ and they are more abundant in healthy carriers than in acute IM patients, predominantly expressing TNFα and IFNγ^46^. Consistent with this, the cytokine response during organoid primary EBV infection was dominated by type I IFN, TNFα, RANTES, and TNFβ, but without significant changes in IFNγ, suggesting a microenvironment resembling asymptomatic infection.

In summary, the tonsil organoid model provides a highly tractable platform to investigate EBV infection-mediated changes in B cell biology and concurrent T cell responses. The presence of infected and uninfected B cell subsets in the same microenvironment also highlights B cell heterogeneity during initial EBV infection as well as the distinction between tonsil organoids and other contemporary *in vitro* platforms. Simultaneous assessment of cellular immune responses to EBV can also readily be modeled in organoids established from seronegative or seropositive donors. The EBV tonsil organoid model offers a versatile platform to investigate EBV pathogenesis, antiviral responses and therapeutic strategies not accessible in conventional animal models or standard *in vitro* systems.

### Methods

#### Tissue processing, organoid preparation and infection with EBV

Tonsil pairs (n=63) from healthy individuals were collected from the Cooperative Human Tissue Network (CHTN) at the Vanderbilt Medical Center (VUMC) or from the University of California Irvine Medical Center (UCIMC; ethics approved by the UCI IRB #2020-6075) from Children’s Hospital of Orange County (CHOC). Patients underwent tonsillectomy for obstructive sleep apnea or hypertrophy. After surgery, the tonsil pairs from CHTN were put in fresh RPMI with antibiotics and, maintaining the cold chain, shipped overnight to the Wagar lab at UCI. Spleen samples (n=4) were collected from patients undergoing splenectomy at the UCIMC and CHOC. Whole blood was collected from some of the tonsil and spleen tissue donors at UCIMC and CHOC. Additionally whole blood (n=5) was also collected from UCI Institute for Clinical and Translational Science (ICTS). After receiving tonsil pairs, they were processed as previously described^34^. In brief, the tonsils were mechanically dissociated in Ham’s F12 media with 2% FBS in a metal processing cup. The cell suspensions were then filtered through a 100 μm nylon strainer and subjected to density gradient centrifugation. The purified live tonsillar cells were enumerated and cryopreserved in FBS with 10% DMSO at −140°C until use. The spleen tissues were processed on the same day post-surgery following the procedure described above. The serum and PBMCs were separated from whole blood samples by centrifuging the whole blood at 1000g for 5 minutes at room temperature. The serum and PBMC samples were stored at -80°C and −140°C respectively until used. The demographic information of all the tissue donors is provided in **Extended Data Table 1**.

To generate the tonsil organoids, cryopreserved aliquots were thawed, enumerated, and plated in transwells or ultra-low attachment (ULA) 96 well flat bottom plates. The final density of cells was 12×10^6^ cell/ml in a final volume of 500uL (6×10^6^ cells per well) in the transwells or 7.5 x 10^6^ cells/ml in a final volume of 200 μL (1.5 x 10^6^ cells per well) in the 96 well ULA plates. Spleen organoids and PMBCs were plated in the 96 well plates having 7.5 x 10^6^ cells/ml in a final volume of 200 μL (1.5 x 10^6^ cells per well). Following plating tonsil cells, splenocytes or PBMCs, GFP-expressing EBV was added in respective wells to generate EBV-infected organoids. The organoid media was composed of RPMI1640 with glutamax, 10% FBS, 1x non-essential amino acids, 1x sodium pyruvate, 1x penicillin/streptomycin, 1x Normocin, 1x insulin/selenium/transferrin cocktail and 1 μg/mL of human BAFF (produced recombinantly and purified in-house or from Biolegend or SinoBiological).

The optimal dose of the GFP-expressing EBV was established based on the ability to detect EBV-infected tonsil B cells by flow cytometry. A MOI of 0.000005, 0.00005 and 0.0005 of the GFP-expressing EBV (AKATA) were used to infect the organoids and cells were harvested after 2 hours, on days 2, 5, 7, 14 post infection and subjected to flow cytometry analysis for detecting GFP^+^ EBV^+^ B cells. MOI of 0.0005 GFP-expressing EBV was found to be sufficient for detecting EBV^+^ B cells in organoids by day 14. For infecting splenocytes and PBMCs the same dose of EBV was used. The cells were incubated at 37°C, 5% CO_2_ and 1000uL or 100μL of fresh organoid media was added every 2-3 days or every other day by replacing approximately 500uL or 100μL media per well for the transwell or the 96 well plates respectively.

#### Cell depletion and organoid preparation

For naive or memory B cell depletion experiments the *ex vivo* tonsil samples were stained with naive B and memory B cell markers and subjected to FACS sorter for a 2-way sort (naive or memory B cells in one FACS tube and everything else in another FACS tube). Following depletion of naive or memory B cells, organoids were prepared from each of the depleted conditions. For the depleted controls, the naive or memory B cells were added back to other non-sorted cells in their native proportions and organoids were prepared. For depleting total CD4 or CD8 T cells we used MACS columns. In short, the cryopreserved tonsil cells are stained with CD4 or CD8 MACS beads following manufacturer’s instructions. LS columns were used to separate the CD4 or CD8 T cells using a positive selection method. After separation, cells were enumerated and organoids prepared from the flow though portions (CD4 or CD8 depleted). For non-depleted controls, CD4 or CD8 T cells were added back to the flow though in their native proportions. The depleted and non-depleted organoids were maintained according to the above-mentioned method.

#### Flow cytometry

For cell surface characterization of the immune cells in the *ex vivo* tonsil tissues and organoids, cells were stained with separate B cell and T cell or a combined panel panel following established staining protocols. Briefly, thawed cells or harvested organoids were washed with FACS buffer (PBS + 0.1% BSA, 0.05% sodium azide and 2mM EDTA) and stained with live-dead staining dye (Zombie aqua) in PBS for 10 minutes on ice. Then cells were washed with FACS buffer and stained with a cocktail of antibodies (Biolegend) in FACS buffer for 30 minutes on ice. The stained cells were washed twice with the FACS buffer and analyzed using either Novocyte Quanteon (ACEA) instrument or the Northern lights or Aurora spectral cytometer and the acquired data was analyzed using FlowJo software (v10.8.1).

#### Serotyping for EBV

For testing prior exposure to EBV, serum samples were tested for anti-VCA IgG and anti-EBNA1 IgG according to manufacturer’s (Abcam) guideline. In brief, the serum samples were diluted 1:100 with IgG sample diluent. 100uL of the diluted samples and standards (IgG positive control, IgG negative control, IgG cut-off control) were added in duplicates in the precoated (VCA or EBNA1) 96 well plate. The plates were sealed and incubated at 37C for an hour. After that the contents of the well were aspirated, washed three times with a wash buffer and Add 100 µL EBV anti-IgG HRP Conjugate were added followed by 30 minutes incubation at room temperature. After washing three times 100uL of TMB substrate was added in each well and incubated for 15 minutes. 100uL stop solution was added and the absorbance was measured at 450 nm. Sero-status for EBV was calculated based on standard units (SU; mean sample absorbancex10/cut-off) i.e. SU> 11 positive, SU<9 negative.

#### Cell sorting and single cell RNA sequencing

For scRNAseq EBV kinetics analyses from EBV-infected or uninfected organoids from days 4, 7, 14, and 21 were harvested and cells were washed with FACS buffer to remove any residual antibodies/factors generated during culture. Samples were stained with a cocktail containing fluorescent-labeled antibodies DNA oligo-tagged antibodies and sample-specific hashtag antibodies (BioLegend). Samples were washed with FACS buffer and on days 4 and 7 total CD45^+^ immune cells were sorted. We also sorted CD45^+^ immune cells from *ex vivo* tonsil samples of the donors. On days 14 and 21 we sorted EBV^+^ B cells (based on GFP expression), nonB cells from infected and uninfected organoids together and uninfected B cells from EBV-infected and uninfected organoids together. Sample hashing with hashtag oligos (HTOs) allowed us to pool several stimulation conditions into one tube. Pooled samples were resuspended in ice-cold PBS with 0.04% BSA in a final concentration of 2500 cells/μl. Single-cell suspensions were then immediately loaded on a 10X Genomics Chromium Controller with a loading target of 30,000 cells. Libraries were generated using the Chromium Next Gem Single Cell 5’ Reagent Kit v2 (Dual Index) per the manufacturer’s instructions, with additional steps for amplification of TotalSeq/HTO barcodes and V(D)J libraries (10X Genomics). Quality and quantity of libraries were measured on tapestation, qubit, and Bioanalyzer, and sequenced on Illumina NovaSeq 6000 with a sequencing target of 1500 million paired reads per cell for gene expression libraries, and 150 million paired reads per cell for TCR, and BCR libraries and 150 million paired reads per feature barcode libraries.

#### 5ʹ gene expression analysis

Raw reads from gene expression, TCR, BCR, and HTO libraries (where applicable) were aligned and quantified using cellranger multi 8.0.1 against GRCh38-2024-A which included the EBV genome (KC207813) and GFP transcript as extra chromosomes. VDJ libraries were aligned against refdata-cellranger-vdj-GRCh38-alts-ensembl-7.1.0. Gene expression matrices were loaded with Seurat v5.0.0, and EBV viral genes and GFP were isolated into a separate expression slot. TCR gene expression was aggregated (TRA, TRB, TRG, TRD) by gene family. HTO classification and demultiplexing were performed via the HTODemux function. A lower threshold of nFeature_RNA > 200 was applied across all samples. Upper thresholds for nFeature_RNA were adjusted based on sample type: a threshold of nFeature_RNA < 4000 was applied to baseline (Day 0) and non–B cell samples, while EBV-enriched B cell samples from Day 14 and Day 21 were filtered with an upper threshold of nFeature_RNA < 6000. Additionally, a mitochondrial content threshold of <10% was applied to exclude cells with elevated mitochondrial gene expression.

Data normalization and variance stabilization were performed on the integrated object using NormalizeData and ScaleData functions in Seurat. Dimension reduction was performed using the RunPCA function and RunUMAP function was used to generate cell clustering at various resolutions. Cell types were broadly assigned as *CD3D* expressing T cells and *CD19* expressing B cells, *CD19-CD3-* (nonBT) cells which were further subsetted for reclustering individually as described above. Gene and protein markers for each cluster were identified using FindMarkers function. For gene expression, we used Wilcoxon Rank Sum tests for differential marker detection using a log_2_ fold change cutoff of at least 0.25 and clusters were identified using a known catalog of markers defined for T and B cells from human tonsil. A similar approach was followed for protein expression, following normalization of antibody capture using centered logratio (CLR) transformation. A combination of both positive and negative protein markers from each cluster was used to aid annotations. Cluster-specific differential expression analysis across conditions was tested using the Wilcoxon rank-sum test with default settings in Seurat. Only statistically significant genes (log_2_ fold change cutoff of 0.4; adjusted p-value < 0.05) were included in downstream analyses.

To identify transcriptional programs in EBV-infected tonsil organoids in T and B cells, we performed topic modeling on scRNA-seq data using the R package fastTopics (v0.6-192)^85^. Cells were subset from previously annotated Seurat objects which included T and B cell subclusters derived from samples collected at days 4, 7, 14, and 21 post-infections. Raw UMI counts were extracted from the RNA assay and filtered to remove genes with zero total counts across cells. Topic modeling was performed with the fit_topic_model function with default parameters. Topic - specific differentially expressed genes were determined with the de_analysis function and filtered to retain genes with a lfsr < 0.01. The top differentially expressed genes in each topic were used to infer the transcriptional program.

#### BCR analysis

The BCR sequence data was processed with Immcantation (https://immcantation.readthedocs.io/). Genes were first assigned with IgBLAST 1.21.0. Next, heavy V gene allele reassignment was performed with TIgGER 1.0.0. Prior to defining clonal families, hamming distances were calculated with SHazaM 1.1.2 distToNearest fucntion for cells with the same V gene, J gene, and junction length. Clonal families were then constructed with SCOPer 1.2.1 hierarchicalClones in single-cell mode. Germline sequences were inferred with Dowser 1.1.0 using sequences from the IMGT database (compiled November 2024). SHM was calculated with the SHazaM observedMutations function. Clonal lineage trees were constructed with heavy chain sequences with IgPhyML 1.1.5 and visualized with ggtree 3.6.0. Statistics were calculated with ggpubr 0.6.0.

#### TCR analysis

TCR sequence data were processed using Immunarch (https://immunarch.com/). Briefly, annotations generated by cellranger were filtered based on cells that passed all filters in the final UMAP. Analysis was performed on a cluster-, time-, stimulation and donor-specific manner. Data were first parsed on Immunarch using the *repLoad* function with a paired option, where only cells with one productive alpha and one productive beta chain were retained for downstream analyses. Clonality (size, top clones, rare clones, and clonal homeostasis) was examined using the *repExplore* function. Diversity estimates (D50 and Chao) were measured using the *repDiversity* function.

### Author Contributions

MTM: Designed and performed experiments, data curation, formal analysis, investigation, methodology, visualization, writing original draft, and writing review & editing. SS, TBY: data analysis, visualization, writing review & editing. YS, ZW, EMJ: experiments and data curation, writing review & editing. AMS: data curation, resources, and writing review & editing. JMK, AN: writing review & editing. AD, NS: data curation. GA, QZ, DT, LCK, MTW, CF: sample procurement. BG: conceptualization, resources, supervision, project administration, writing review & editing. LEW: conceptualization, data curation, funding acquisition, investigation, project administration, supervision, visualization, and writing original draft and review & editing.

## Acknowledgements

We thank the tonsillectomy patients for their participation in this study; the UCI Institute for Immunology Flow Cytometry Core for technical assistance; Dr. Eric Pearlman for equipment access; Dr. Huw Davies and Dr. Jenny Davies for critical review of the manuscript; UCI Genomics Research and Technology Hub (GRT-Hub) for assistance with sequencing the 10x libraries, Drs. Edwards and Tifrea at UCI Medical Center for coordinating sample collection; Dr. Saligrama and the Genome Technology Access Center at the McDonnell Genome Institute at Washington University School of Medicine for help with sequencing the 10x libraries and the Cooperative Human Tissue Network (CHTN) for providing deidentified pediatric tonsil samples. The CHTN consists of six academic institutions that collect and distribute remnant human biospecimens from routine surgical and autopsy procedures to support biomedical research.

This work was supported by funding from the Wellcome Leap HOPE Program (to L.E.W.), the National Institutes of Health (NIH) grant R01AI173023 (to L.E.W.), by R01CA228700 and R01DE033907 to B.E.G, by an American Cancer Society Discovery Boost Grant to B.E.G, by the George and Sandra K. Schussel endowed chair to B.E.G., by Immunology Training grant GU12093 to M.T.M., and by American Cancer Society Post-doctoral Fellowship PF-23-1144614-01-IBCD to Y.S. Select sample collections were supported by the National Center for Advancing Translational Sciences, National Institutes of Health, through Grant UM1 TR004927. The content is solely the responsibility of the authors and does not necessarily represent the official views of the funders.

## Lead contact

Further information and requests for resources and reagents should be directed to and will be fulfilled by the lead contacts, Lisa Wagar and Benjamin Gewurz.

## Data and code availability

All single-cell RNA-seq data reported in this paper will be uploaded to GEO prior to formal acceptance of this manuscript. Any additional information required to reanalyze the data reported in this paper is available from the lead contact upon request.

## REFERENCES

1. Damania, B., Kenney, S. C. & Raab-Traub, N. Epstein-Barr virus: Biology and clinical disease. Cell 185, 3652–3670 (2022).

2. Cohen, J. I., Fauci, A. S., Varmus, H. & Nabel, G. J. Epstein-Barr Virus: An Important Vaccine Target for Cancer Prevention. Sci. Transl. Med. 3, 107fs7–107fs7 (2011).

3. Wong, Y., Meehan, M. T., Burrows, S. R., Doolan, D. L. & Miles, J. J. Estimating the global burden of Epstein-Barr virus-related cancers. J. Cancer Res. Clin. Oncol. 148, 31–46 (2022).

4. Robinson, W. H., Younis, S., Love, Z. Z., Steinman, L. & Lanz, T. V. Epstein–Barr virus as a potentiator of autoimmune diseases. Nat. Rev. Rheumatol. 20, 729–740 (2024).

5. Luzuriaga, K. & Sullivan, J. L. Infectious Mononucleosis. N. Engl. J. Med. 362, 1993–2000 (2010).

6. Thorley-Lawson, D. A. EBV Persistence—Introducing the Virus. in Epstein Barr Virus Volume 1 (ed. Münz, C.) vol. 390 151–209 (Springer International Publishing, Cham, 2015).

7. Yu, H. & Robertson, E. S. Epstein–Barr Virus History and Pathogenesis. Viruses 15, 714 (2023).

8. Babcock, G. J., Decker, L. L., Volk, M. & Thorley-Lawson, D. A. EBV Persistence in Memory B Cells In Vivo. Immunity 9, 395–404 (1998).

9. Thorley-Lawson, D. A., Hawkins, J. B., Tracy, S. I. & Shapiro, M. The pathogenesis of Epstein–Barr virus persistent infection. Curr. Opin. Virol. 3, 227–232 (2013).

10. Guo, R. & Gewurz, B. E. Epigenetic control of the Epstein-Barr lifecycle. Curr. Opin. Virol. 52, 78–88 (2022).

11. Münz, C. Epstein–Barr virus pathogenesis and emerging control strategies. Nat. Rev. Microbiol. 1–13 (2025) doi:10.1038/s41579-025-01181-y.

12. Chiu, Y.-F., Ponlachantra, K. & Sugden, B. How Epstein Barr Virus Causes Lymphomas. Viruses 16, 1744 (2024).

13. Price, A. M. & Luftig, M. A. To Be or Not IIb: A Multi-Step Process for Epstein-Barr Virus Latency Establishment and Consequences for B Cell Tumorigenesis. PLoS Pathog. 11, e1004656 (2015).

14. Murata, T. et al. Molecular Basis of Epstein–Barr Virus Latency Establishment and Lytic Reactivation. Viruses 13, 2344 (2021).

15. Babcock, G. J., Hochberg, D. & Thorley-Lawson, D. A. The Expression Pattern of Epstein-Barr Virus Latent Genes In Vivo Is Dependent upon the Differentiation Stage of the Infected B Cell. Immunity 13, 497–506 (2000).

16. Joseph, A. M., Babcock, G. J. & Thorley-Lawson, D. A. Cells Expressing the Epstein-Barr Virus Growth Program Are Present in and Restricted to the Naive B-Cell Subset of Healthy Tonsils. J. Virol. 74, 9964–9971 (2000).

17. Roughan, J. E. & Thorley-Lawson, D. A. The Intersection of Epstein-Barr Virus with the Germinal Center. J. Virol. 83, 3968–3976 (2009).

18. Roughan, J. E., Torgbor, C. & Thorley-Lawson, D. A. Germinal Center B Cells Latently Infected with Epstein-Barr Virus Proliferate Extensively but Do Not Increase in Number. J. Virol. 84, 1158–1168 (2010).

19. Souza, T. A., Stollar, B. D., Sullivan, J. L., Luzuriaga, K. & Thorley-Lawson, D. A. Peripheral B cells latently infected with Epstein–Barr virus display molecular hallmarks of classical antigen-selected memory B cells. Proc. Natl. Acad. Sci. U. S. A. 102, 18093–18098 (2005).

20. Kurth, J., Hansmann, M.-L., Rajewsky, K. & Küppers, R. Epstein–Barr virus-infected B cells expanding in germinal centers of infectious mononucleosis patients do not participate in the germinal center reaction. Proc. Natl. Acad. Sci. U. S. A. 100, 4730–4735 (2003).

21. SoRelle, E. D. et al. Time-resolved transcriptomes reveal diverse B cell fate trajectories in the early response to Epstein-Barr virus infection. Cell Rep. 40, 111286 (2022).

22. Taylor, G. S., Long, H. M., Brooks, J. M., Rickinson, A. B. & Hislop, A. D. The Immunology of Epstein-Barr Virus–Induced Disease. Annu. Rev. Immunol. 33, 787–821 (2015).

23. Chijioke, O., Landtwing, V. & Münz, C. NK Cell Influence on the Outcome of Primary Epstein–Barr Virus Infection. Front. Immunol. 7, (2016).

24. Long, H. M., Meckiff, B. J. & Taylor, G. S. The T-cell Response to Epstein-Barr Virus–New Tricks From an Old Dog. Front. Immunol. 10, (2019).

25. Liu, M., Wang, R. & Xie, Z. T cell-mediated immunity during Epstein–Barr virus infections in children. Infect. Genet. Evol. 112, 105443 (2023).

26. Lelic, A. et al. The Polyfunctionality of Human Memory CD8+ T Cells Elicited by Acute and Chronic Virus Infections Is Not Influenced by Age. PLOS Pathog. 8, e1003076 (2012).

27. Lam, J. K. P. et al. Emergence of CD4+ and CD8+ Polyfunctional T Cell Responses Against Immunodominant Lytic and Latent EBV Antigens in Children With Primary EBV Infection. Front. Microbiol. 9, 416 (2018).

28. Berhan, A., Bayleyegn, B. & Getaneh, Z. HIV/AIDS Associated Lymphoma: Review. Blood Lymphat. Cancer Targets Ther. 12, 31–45 (2022).

29. Zhong, L. et al. Urgency and necessity of Epstein-Barr virus prophylactic vaccines. Npj Vaccines 7, 1–14 (2022).

30. Münz, C. EBV Infection and Its Immune Control in Humanized Mice. in 1–23 (Springer, Berlin, Heidelberg). doi:10.1007/82_2025_285.

31. Münz, C. Recent advances in animal models of lymphomagenesis caused by human γ-herpesviruses. Curr. Opin. Virol. 71, 101461 (2025).

32. Ma, S.-D. et al. A New Model of Epstein-Barr Virus Infection Reveals an Important Role for Early Lytic Viral Protein Expression in the Development of Lymphomas. J. Virol. 85, 165–177 (2011).

33. Ahmed, E. H. & Baiocchi, R. A. Murine Models of Epstein-Barr Virus–Associated Lymphomagenesis. ILAR J. 57, 55–62 (2016).

34. Wagar, L. E. et al. Modeling human adaptive immune responses with tonsil organoids. Nat. Med. 27, 125–135 (2021).

35. Kastenschmidt, J. M. et al. Influenza vaccine format mediates distinct cellular and antibody responses in human immune organoids. Immunity 56, 1910–1926.e7 (2023).

36. Mitul, M. T. et al. Tissue-specific sex differences in pediatric and adult immune cell composition and function. Front. Immunol. 15, (2024).

37. Wagoner, Z. W. et al. Systems immunology analysis of human immune organoids identifies host-specific correlates of protection to different influenza vaccines. Cell Stem Cell 32, 529–546.e6 (2025).

38. Sureshchandra, S. et al. Deep profiling of human T cells defines compartmentalized clones and phenotypic trajectories across blood and tonsils. Immunity 58, 3130–3143.e8 (2025).

39. Kanda, T., Yajima, M., Ahsan, N., Tanaka, M. & Takada, K. Production of High-Titer Epstein-Barr Virus Recombinants Derived from Akata Cells by Using a Bacterial Artificial Chromosome System. J. Virol. 78, 7004–7015 (2004).

40. Dorner, M. et al. Distinct Ex Vivo Susceptibility of B-Cell Subsets to Epstein-Barr Virus Infection According to Differentiation Status and Tissue Originᰔ. J VIROL 82, 13 (2008).

41. Chaganti, S. et al. Epstein-Barr virus colonization of tonsillar and peripheral blood B-cell subsets in primary infection and persistence. Blood 113, 6372–6381 (2009).

42. Heath, E. et al. Epstein-Barr Virus Infection of Naïve B Cells In Vitro Frequently Selects Clones with Mutated Immunoglobulin Genotypes: Implications for Virus Biology. PLoS Pathog. 8, e1002697 (2012).

43. Victora, G. D. & Nussenzweig, M. C. Germinal Centers. Annu. Rev. Immunol. 40, 413–442 (2022).

44. Elsner, R. A. & Shlomchik, M. J. Germinal Center and Extrafollicular B Cell Responses in Vaccination, Immunity, and Autoimmunity. Immunity 53, 1136–1150 (2020).

45. Precopio, M. L., Sullivan, J. L., Willard, C., Somasundaran, M. & Luzuriaga, K. Differential Kinetics and Specificity of EBV-Specific CD4+ and CD8+ T Cells During Primary Infection1. J. Immunol. 170, 2590–2598 (2003).

46. Meckiff, B. J. et al. Primary EBV Infection Induces an Acute Wave of Activated Antigen-Specific Cytotoxic CD4+ T Cells. J. Immunol. Baltim. Md 1950 203, 1276–1287 (2019).

47. Tanner, J., Weis, J., Fearon, D., Whang, Y. & Kieff, E. Epstein-Barr virus gp350/220 binding to the B lymphocyte C3d receptor mediates adsorption, capping, and endocytosis. Cell 50, 203–213 (1987).

48. Robson, S. C., Sévigny, J. & Zimmermann, H. The E-NTPDase family of ectonucleotidases: Structure function relationships and pathophysiological significance. Purinergic Signal. 2, 409 (2006).

49. Regateiro, F. S., Cobbold, S. P. & Waldmann, H. CD73 and adenosine generation in the creation of regulatory microenvironments. Clin. Exp. Immunol. 171, 1–7 (2013).

50. Figueiró, F. et al. Phenotypic and functional characteristics of CD39 ^high^ human regulatory B cells (Breg). OncoImmunology 5, e1082703 (2016).

51. Grimm, J. M., et al. Prospective studies of infectious mononucleosis in university students. Clin. Transl. Immunol. 5, e94 (2016).

52. Crawford, D. H., et al. A cohort study among university students: identification of risk factors for Epstein-Barr virus seroconversion and infectious mononucleosis. Clin. Infect. Dis. Off. Publ. Infect. Dis. Soc. Am. 43, 276–282 (2006).

53. Silins, S. L. et al. Asymptomatic primary Epstein-Barr virus infection occurs in the absence of blood T-cell repertoire perturbations despite high levels of systemic viral load. Blood 98, 3739–3744 (2001).

54. Dunmire, S. K., Verghese, P. S. & Balfour, H. H. Primary Epstein-Barr virus infection. J. Clin. Virol. 102, 84–92 (2018).

55. Martín-Pérez, D. et al. Epstein-Barr virus microRNAs repress BCL6 expression in diffuse large B-cell lymphoma. Leukemia 26, 180–183 (2012).

56. Pei, Y., Banerjee, S., Jha, H. C., Sun, Z. & Robertson, E. S. An essential EBV latent antigen 3C binds Bcl6 for targeted degradation and cell proliferation. PLOS Pathog. 13, e1006500 (2017).

57. Dey, K. K., Hsiao, C. J. & Stephens, M. Visualizing the structure of RNA-seq expression data using grade of membership models. PLOS Genet. 13, e1006599 (2017).

58. Carbonetto, P. et al. GoM DE: interpreting structure in sequence count data with differential expression analysis allowing for grades of membership. *bioRxiv* 2023.03.03.531029 (2023) doi:10.1101/2023.03.03.531029.

59. SoRelle, E. D., Reinoso-Vizcaino, N. M., Horn, G. Q. & Luftig, M. A. Epstein-Barr virus perpetuates B cell germinal center dynamics and generation of autoimmune-associated phenotypes in vitro. Front. Immunol. 13, (2022).

60. Choi, I.-K. et al. Signaling by the Epstein-Barr virus LMP1 protein induces potent cytotoxic CD4+ and CD8+ T cell responses. Proc. Natl. Acad. Sci. U. S. A. 115, E686–E695 (2018).

61. Brooks, J. M. et al. Early T Cell Recognition of B Cells following Epstein-Barr Virus Infection: Identifying Potential Targets for Prophylactic Vaccination. PLOS Pathog. 12, e1005549 (2016).

62. Torgbor, C., Thorley-Lawson, D. A. & Moormann, A. M. Epstein-Barr virus-infected tonsillar marginal zone B cells in vivo as a precursor for immunosuppression-related B-cell lymphoma. J. Virol. 99, e01051–24 (2025).

63. Boccellato, F. et al. EBNA2 Interferes with the Germinal Center Phenotype by Downregulating BCL6 and TCL1 in Non-Hodgkin’s Lymphoma Cells. J. Virol. 81, 2274–2282 (2007).

64. Sun, K., Bose, D., Singh, R. K., Pei, Y. & Robertson, E. S. The F-box E3 ligase protein FBXO11 regulates EBNA3C-associated degradation of BCL6. J. Virol. 98, e00548–24.

65. Riel, M. & Johannsen, E. C. LMP2A—The Other EBV Oncogene. in 1–37 (Springer, Berlin, Heidelberg). doi:10.1007/82_2025_330.

66. Cen, O. & Longnecker, R. Latent Membrane Protein 2 (LMP2). in Epstein Barr Virus Volume 2: One Herpes Virus: Many Diseases (ed. Münz, C.) 151–180 (Springer International Publishing, Cham, 2015). doi:10.1007/978-3-319-22834-1_5.

67. Kieser, A. The Latent Membrane Protein 1 (LMP1): Biological Functions and Molecular Mechanisms. Curr. Top. Microbiol. Immunol. 10.1007/82_2025_321 (2025) doi:10.1007/82_2025_321.

68. Dai, J. et al. Epstein-Barr virus induces germinal center light zone chromatin architecture and promotes survival through enhancer looping at the BCL2A1 locus. mBio 15, e02444–23 (2023).

69. Epeldegui, M., Hung, Y. P., McQuay, A., Ambinder, R. F. & Martínez-Maza, O. Infection of human B cells with Epstein-Barr virus results in the expression of somatic hypermutation-inducing molecules and in the accrual of oncogene mutations. Mol. Immunol. 44, 934–942 (2007).

70. Wang, L. W. et al. Epstein-Barr-Virus-Induced One-Carbon Metabolism Drives B Cell Transformation. Cell Metab. 30, 539–555.e11 (2019).

71. Souza, T. A., Stollar, B. D., Sullivan, J. L., Luzuriaga, K. & Thorley-Lawson, D. A. Influence of EBV on the Peripheral Blood Memory B Cell Compartment1. J. Immunol. 179, 3153–3160 (2007).

72. He, B., Raab-Traub, N., Casali, P. & Cerutti, A. EBV-Encoded Latent Membrane Protein 1 Cooperates with BAFF/BLyS and APRIL to Induce T Cell-Independent Ig Heavy Chain Class Switching. J. Immunol. Baltim. Md 1950 171, 5215–5224 (2003).

73. Uchida, J. et al. Mimicry of CD40 Signals by Epstein-Barr Virus LMP1 in B Lymphocyte Responses. Science 286, 300–303 (1999).

74. Zhang, B. et al. Immune Surveillance and Therapy of Lymphomas Driven by Epstein-Barr Virus Protein LMP1 in a Mouse Model. Cell 148, 739–751 (2012).

75. Rastelli, J. et al. LMP1 signaling can replace CD40 signaling in B cells in vivo and has unique features of inducing class-switch recombination to IgG1. Blood 111, 1448–1455 (2008).

76. Stavnezer, J., Guikema, J. E. J. & Schrader, C. E. Mechanism and Regulation of Class Switch Recombination. Annu. Rev. Immunol. 26, 261–292 (2008).

77. Hsu, D. H. et al. Expression of interleukin-10 activity by Epstein-Barr virus protein BCRF1. Science 250, 830–832 (1990).

78. Long, H. M. et al. MHC II tetramers visualize human CD4+ T cell responses to Epstein–Barr virus infection and demonstrate atypical kinetics of the nuclear antigen EBNA1 response. J. Exp. Med. 210, 933–949 (2013).

79. Pudney, V. A., Leese, A. M., Rickinson, A. B. & Hislop, A. D. CD8+ immunodominance among Epstein-Barr virus lytic cycle antigens directly reflects the efficiency of antigen presentation in lytically infected cells. J. Exp. Med. 201, 349–360 (2005).

80. Callan, M. F. C. et al. Large clonal expansions of CD8+ T cells in acute infectious mononucleosis. Nat. Med. 2, 906–911 (1996).

81. Scherrenburg, J., Piriou, E. R. W. A. N., Nanlohy, N. M. & van Baarle, D. Detailed analysis of Epstein–Barr virus-specific CD4+ and CD8+ T cell responses during infectious mononucleosis. Clin. Exp. Immunol. 153, 231–239 (2008).

82. Amarillo, M. E. et al. Tonsillar cytotoxic CD4 T cells are involved in the control of EBV primary infection in children. Sci. Rep. 14, 2135 (2024).

83. Barros, M. H. M., Vera-Lozada, G., Segges, P., Hassan, R. & Niedobitek, G. Revisiting the tissue microenvironment of infectious mononucleosis: Identification of EBV infection in T cells and deep characterization of immune profiles. Front. Immunol. 10, (2019).

84. Ning, R. J., Xu, X. Q., Chan, K. H. & Chiang, A. K. S. Long-term carriers generate Epstein–Barr virus (EBV)-specific CD4+ and CD8+ polyfunctional T-cell responses which show immunodominance hierarchies of EBV proteins. Immunology 134, 161–171 (2011).

85. Carbonetto, P. et al. GoM DE: interpreting structure in sequence count data with differential expression analysis allowing for grades of membership. Genome Biol. 24, 236 (2023).

