## Extended Data Figure 1 for "A tonsil organoid model reveals Epstein-Barr virus infected germinal center B cell states during primary infection"

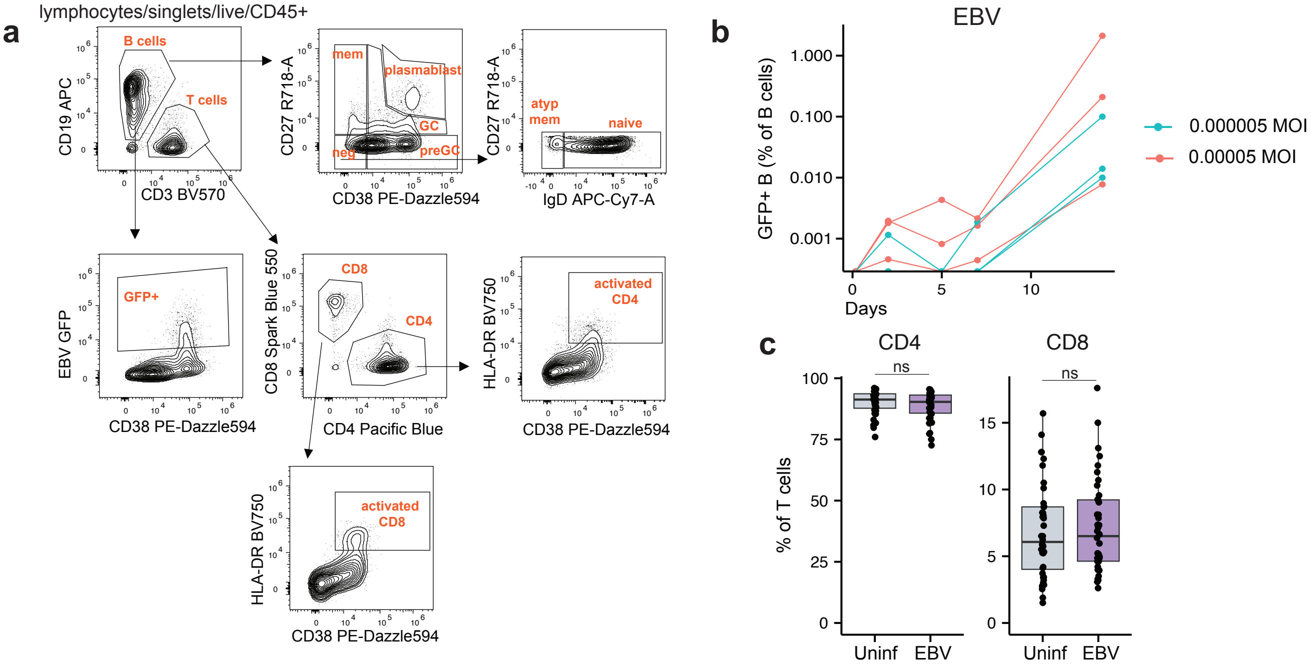


**Extended Data Figure 1: Extended analysis of B- and T-cell responses in tonsil organoids following EBV infection.** (a) Gating strategy: Representative gating strategy for identifying B and T cell subsets in the tonsil organoids. (b) Frequencies of $EBV^{+}$ B cells in response to 0.000005 and 0.00005 MOI of GFP-expressing EBV (n=3). (c) Frequencies of total CD4 ad CD8 T cells in uninfected and EBV-infected tonsil organoids on day 14 (n=40 per group). Mann Whitney U tests were used to calculate p values between groups. Boxplots indicate the median value, with hinges denoting the first and third quartiles and whiskers denoting the highest and lowest value within 1.5 times the interquartile range of the hinges. MOI, multiplicity of infection; ns, not significant ($p \geq 0.05$).
