## Extended Data Figure 2 for "A tonsil organoid model reveals Epstein-Barr virus infected germinal center B cell states during primary infection"

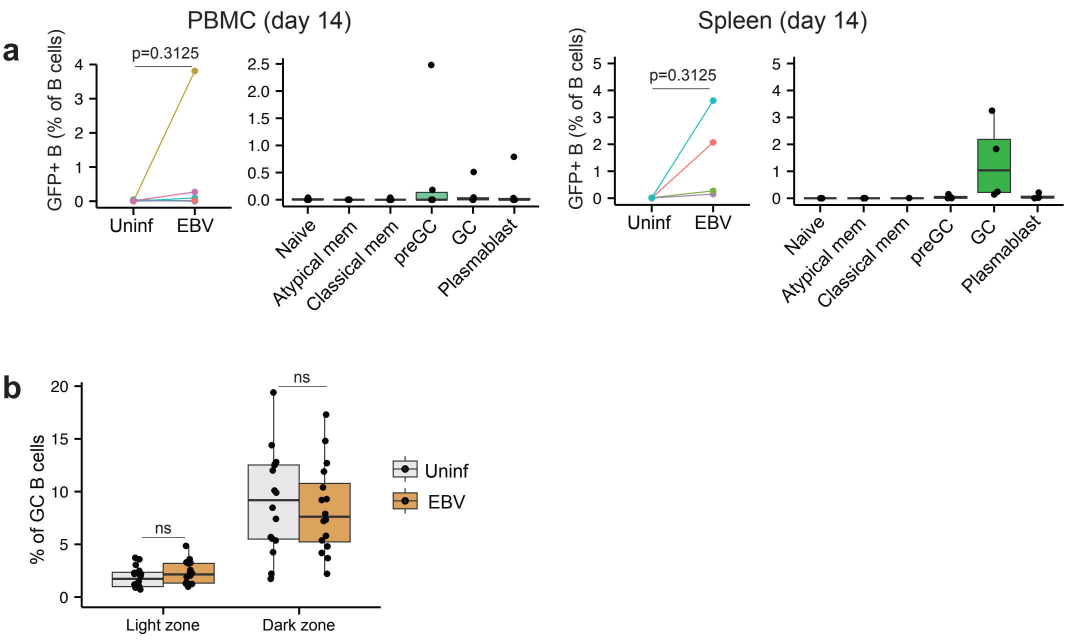


**Extended Data Figure 2: Additional phenotypic differences between EBV$^{+}$ and EBV$^{-}$ B cells.** (a) EBV$^{+}$ B cell frequencies and phenotypes on day 14 in uninfected and EBV-infected PBMCs (n=6 per group) and spleen organoids (n=4 per group). (b) Light zone (CD83$^{+}$) and dark zone (CXCR4$^{+}$) GC B cell frequencies in the uninfected and EBV-infected tonsil organoids on day14 (n=16 per group). Mann Whitney U tests were used to calculate p values between groups. Boxplots indicate the median value, with hinges denoting the first and third quartiles and whiskers denoting the highest and lowest value within 1.5 times the interquartile range of the hinges. ns, not significant ($p \geq 0.05$).
