## Extended Data Figure 3 for "A tonsil organoid model reveals Epstein-Barr virus infected germinal center B cell states during primary infection"

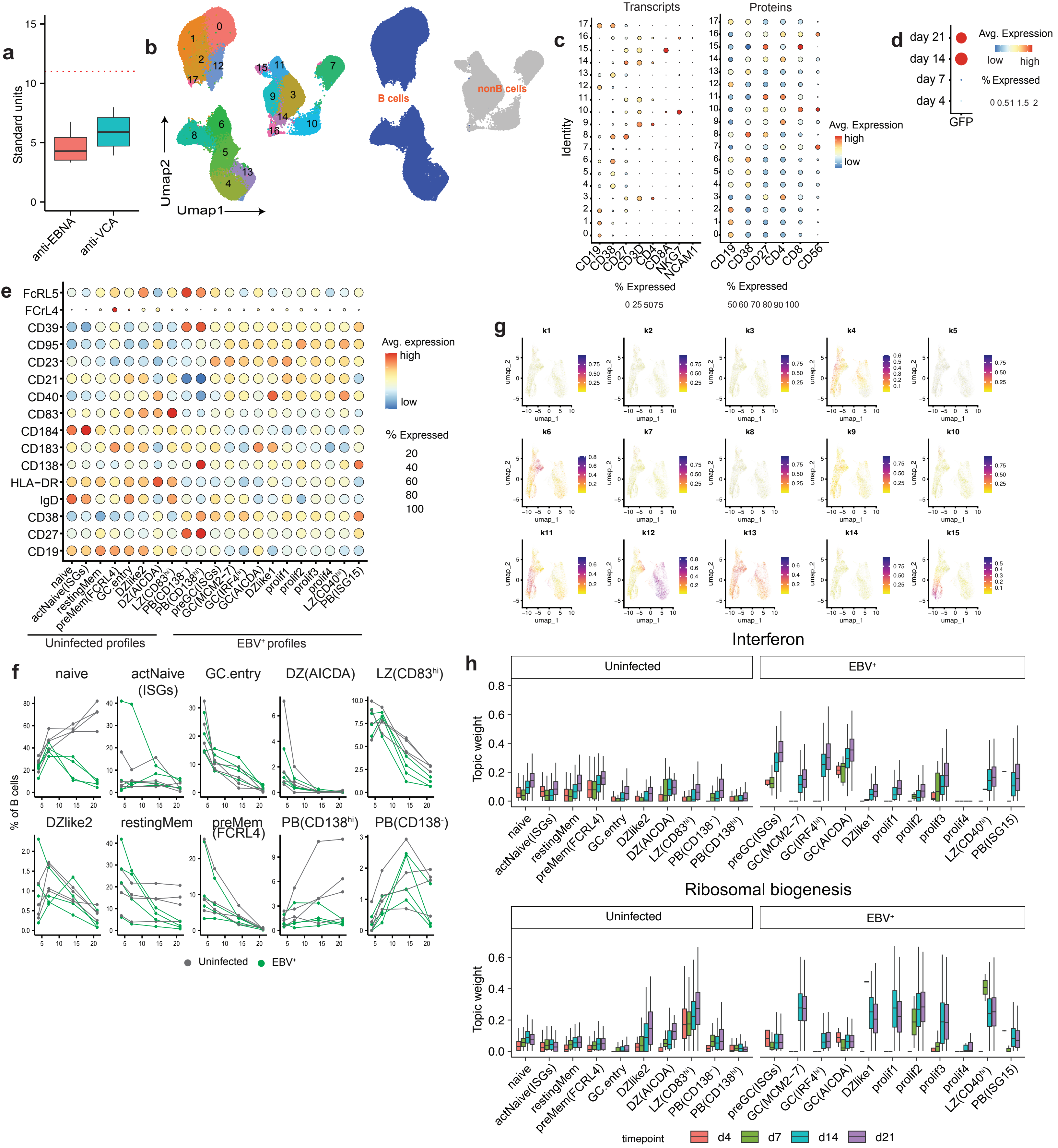

**Extended Data Figure 3: Features of EBV⁺ B cells in primary infection.** (a) anti-EBNA and anti-VCA antibodies in donors used in the scRNAseq experiment (n=4). The red dotted line indicates the baseline for seropositivity. (b, c) UMAP and features of B and nonB cells sorted from uninfected and EBV-infected organoids (89,444 cells, all donors, timepoint and stimulation combined, n=4, 1 experiment). (d) Detection of GFP transcripts in infected organoids across time. (e) Bubble plot of cell surface markers used to identify B cell clusters. Bubble size indicates percentage of cells expressing the marker and color indicates magnitude of expression. (f) B cells cluster frequencies in EBV-infected and uninfected organoids from day 4 to day 21 (n=4 per group). None of the B cell cluster frequencies were significantly different.(g) Feature plot showing the distribution of topics identified in EBV⁺ and bystander B cells. (h) Interferon and ribosomal biogenesis topic weights in B cell clusters in the EBV-infected organoids across time. Boxplots indicate the median value, with hinges denoting the first and third quartiles and whiskers denoting the highest and lowest value within 1.5 times the interquartile range of the hinges. Mann Whitney U tests followed by multiple hypothesis correction using the Benjamini & Hochberg method were used to calculate p values.
