## Extended Data Figure 4 for "A tonsil organoid model reveals Epstein-Barr virus infected germinal center B cell states during primary infection"

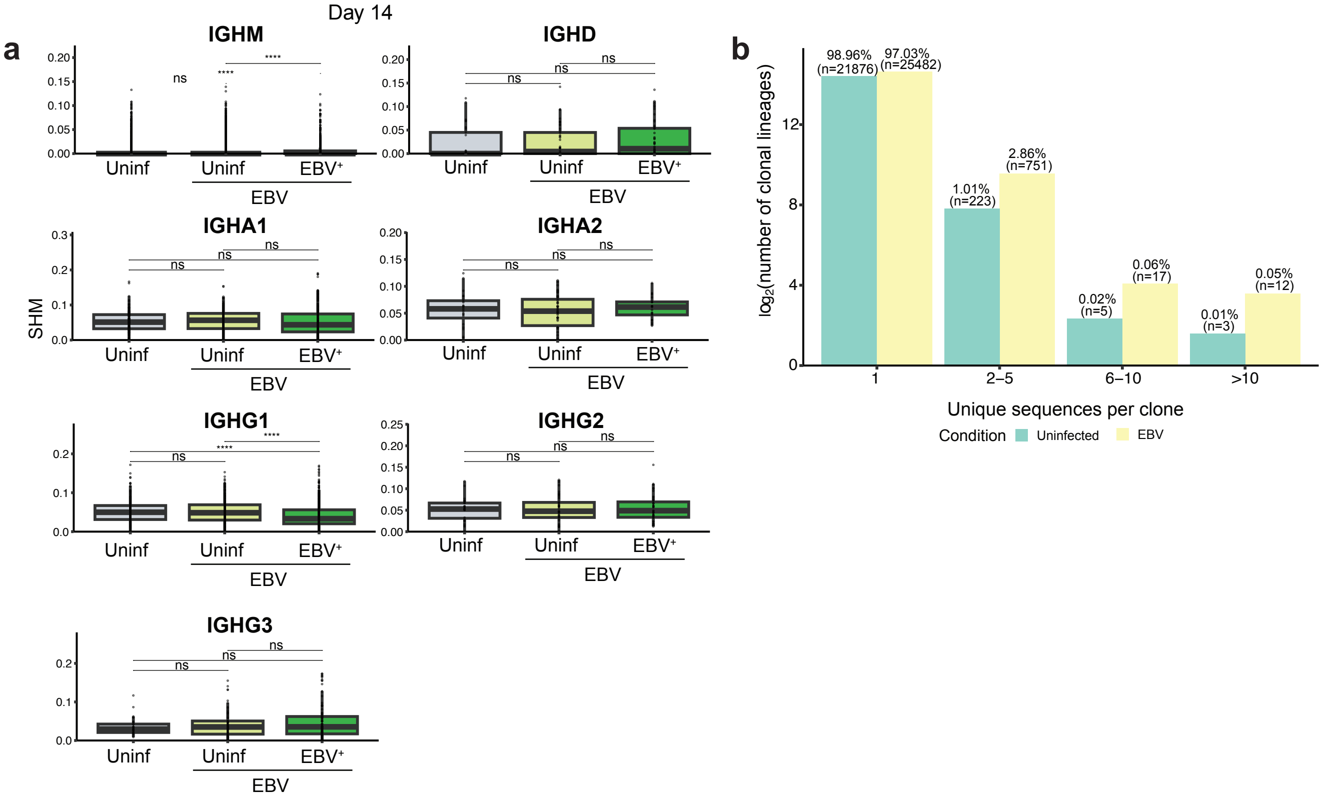


**Extended Data Figure 4: Additional B cell clonal features in EBV-infected tonsil organoids.** (**a**) SHM frequency among immunoglobulin Isotypes in uninfected, EBV-uninfected and EBV⁺ B cells on day 14. (**b**) Clone counts representing unique sequences in uninfected and EBV-infected organoids. Mann-Whitney U tests (two-sided) followed by Benjamini & Hochberg method were used to calculate significance values (*p< 0.05, **p< 0.01, ***p< 0.001, ****p< 0.0001). SHM, Somatic hypermutation; ns, not significant (p ≥ 0.05).
