## Extended Data Figure 5 for "A tonsil organoid model reveals Epstein-Barr virus infected germinal center B cell states during primary infection"

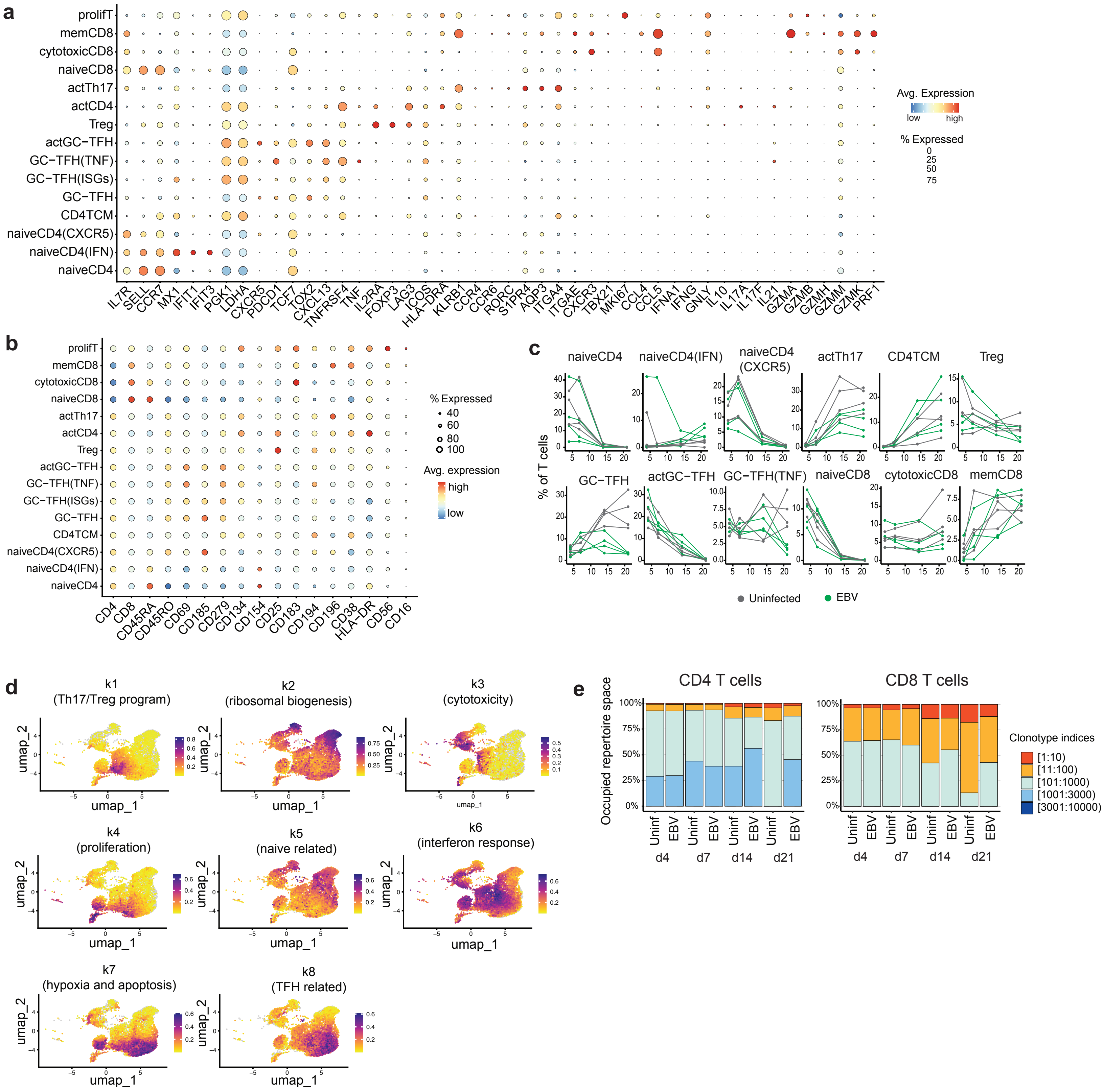


**Extended Data Figure 5: Additional features of T cell response in tonsil organoids in primary EBV infection.** Bubble plots showing the defining (a) transcripts (b) cell surface proteins expressed in T cell subsets in primary EBV infection in the organoids (all donors, timepoints and stimulation aggregated, n=4, 1 experiment). Bubble size indicates percentage of cells expressing the transcript/marker and color indicates magnitude of expression. (c) Frequencies of CD4 T cell clusters in uninfected (gray) and EBV-infected (green) organoids over the time course in primary EBV infection (n=4 per group). None of the CD4 T cell cluster frequencies were signicantly different between uninfected and EBV-infected organoids accros timepoints (d) Feature plot showing the distribution of topics (gene expression program) in T cells (all timepoints, stimulation and donor aggregated). (e) Bar graph representing top clonal proportions of CD4 and CD8 T cells in the uninfected and EBV-infected organoids on 4, 7, 14 and 21 days post infection. Mann Whitney U tests followed by multiple hypothesis correction using the Benjamini & Hochberg method were used to calculate p values.
